# Retrieving chromatin patterns from deep sequencing data using correlation functions

**DOI:** 10.1101/054049

**Authors:** Jana Molitor, Jan-Philipp Mallm, Karsten Rippe, Fabian Erdel

## Abstract

Epigenetic modifications and other chromatin features partition the genome on multiple length scales. They define chromatin domains with distinct biological functions that come in sizes ranging from single modified DNA bases to several megabases in case of heterochromatic histone modifications. Due to chromatin folding, domains that are well separated along the linear nucleosome chain can form long-range interactions in three-dimensional space. It has now become a routine task to map epigenetic marks and chromatin structure by deep sequencing methods. However, assessing and comparing the properties of chromatin domains and their positional relationships across data sets without *a priori* assumptions remains challenging. Here, we introduce multi-scale correlation evaluation (MCORE), which uses the fluctuation spectrum of mapped sequencing reads to quantify and compare chromatin patterns over a broad range of length scales in a model-independent manner. We applied MCORE to map the chromatin landscape in mouse embryonic stem cells and differentiated neural cells. We integrated sequencing data from chromatin immunoprecipitation, RNA expression, DNA methylation and chromosome conformation capture experiments into network models that reflect the positional relationships among these features on different genomic scales. Furthermore, we used MCORE to compare our experimental data to models for heterochromatin reorganization during differentiation. The application of correlation functions to deep sequencing data complements current evaluation schemes and will support the development of quantitative descriptions of chromatin networks.

## Introduction

Most processes in eukaryotic cells that involve interactions with the genome are controlled by the chromatin context. Accordingly, DNA replication, DNA repair, RNA expression and RNA splicing have been found to be regulated by different combinations of DNA methylation (5mC) and histone modifications (1, 2). The genome-wide distribution of these and other chromatin features, like binding sites of transcription factors, contact frequencies between genomic loci and transcriptional activity, can routinely be assessed by deep sequencing (1). Recent methodological developments enable the analysis of low cell numbers or even single cells (3-5), the simultaneous readout of various features (6), and the measurement of site-specific binding dynamics (7). Thus, sequencing data at unprecedented resolution and throughput are becoming available, providing a rich source of information on molecular networks that shape the chromatin landscape. However, there is a gap between the widely used techniques for the qualitative analysis of sequencing data and what is needed for testing biophysical models that quantitatively describe the dynamics of chromatin states and long-range gene regulation (8). Specific objectives are for example to relate the size and shape of modified domains to the underlying formation mechanism, to assess the contribution of chromatin contacts to the establishment and maintenance of chromatin states, and to describe the positional relationship among different marks, which is an important step towards understanding the function of distal regulatory elements.

Currently, deep sequencing data are mostly analyzed on the basis of local enrichments of read density, with the goal to identify regions scoring positive for one or more features of interest. Most of these approaches (see **Table S1** for an incomplete list) fall into two categories, namely peak calling algorithms (9-11) and probabilistic network models (12-14). Identification of enriched regions typically involves assumptions about their characteristic width and enrichment level, and regions above a certain significance level are considered positive. While this strategy is suitable for finding the most highly enriched genomic regions, it does not preserve the information content of complex patterns that involve different enrichment levels and are incompatible with binarization (**Fig. S1**). Furthermore, undersampling, noise and technical bias represent complications that can change the apparent read density at individual loci, thereby introducing or masking similarities between data sets when comparing them based on sets of local enrichments (15-17). Due to these difficulties, peak calling results depend on user-defined input parameters and the specific algorithm used (18, 19). In turn, chromatin state annotations differ with respect to state number, state identity and spatial extension of the corresponding chromatin domains (12, 13). These uncertainties are particularly critical for the study of heterochromatic regions, which contain a combination of broadly distributed histone marks, 5mC and associated proteins (20, 21). Accordingly, quantitative comparisons between the genome-wide topology of heterochromatin domains and the predictions from mechanistic models for the formation and maintenance of heterochromatin states (e.g. (22-24) and references therein) are currently fraught with difficulties.

Here, we introduce an approach termed multi-scale correlation evaluation (MCORE) that complements the above-mentioned repertoire of analysis methods for deep sequencing data. MCORE avoids assumptions about the shape and the amplitude of enriched regions and evaluates all mapped sequencing reads without filtering. It retrieves information from correlation functions, which are used for the discovery of patterns in noisy and possibly undersampled data sets in many fields of research (25-29). The use of correlation functions in the context of deep sequencing has mostly been restricted to strand cross correlation for measuring fragment lengths (18, 30) and short-range autocorrelation for comparing ChIP-seq data sets to each other (31). Key advantages of correlation functions are the intrinsic removal of (white) noise, robust identification of characteristic length scales and straightforward assessment of spatial relationships between two different features. Conveniently, correlation functions can be used to retrieve information about patterns with unknown geometry (**Fig. S1**). We used MCORE to analyze the chromatin landscape of embryonic stem cells (ESCs) and neural cells (neural progenitor/brain cells, NCs) as their differentiated counterparts, focusing on 11 different chromatin features (**Table S2**). These data sets covered histone modifications, DNA methylation, RNA expression, genome folding and binding of chromatin-associated proteins. For each feature we identified the associated nucleosome repeat length and the characteristic domain sizes along with their relative abundance in the genome. In a pair-wise analysis we determined the (anti-)colocalization and positional relationship between features on different genomic scales and used the results to construct network models for chromatin signaling. We compared ESCs to NCs to retrieve information about the spatial reorganization of chromatin during differentiation and to map the global transitions that occurred at active and repressive chromatin domains. Alterations were most pronounced for heterochromatic H3K9me3/H3K27me3 regions that changed their size, their location within chromosome territories and their positioning relative to DNA methylation and to each other.

## Materials and Methods

### Calculation of normalized occupancy profiles

Sequencing reads were mapped to the mouse mm9 assembly using Bowtie (32). Only uniquely mapping hits without mismatches were considered and duplicates were removed. Mapped reads were processed according to the following steps: Bisulfite sequencing (BS-seq) data, which are used to map DNA methylation at single base pair resolution, are usually available as methylation scores calculated from the ratio of converted reads divided by the sum of converted and unconverted reads at a given position. These can be directly used for computing the correlation function as described below. For all other sequencing readouts, the coverage was initially calculated for each chromosome by extending the reads to fragment length, yielding a histogram with the genomic coordinate on the x-axis and the number of reads per base pair on the у-axis. For Hi-C and ChIA-PET data only inter-chromosomal reads were considered to identify the surface of chromosome territories. To calculate normalized occupancy profiles, samples were processed depending on the type of experiment. In general, it is important to account for fragmentation bias, library preparation bias and genome mappability. These multiplicative biases are also included in the input sample and should cancel out in the ratio of specific signal *A* and input signal *I* (*A/I*). In RNA-seq experiments the input signal can be replaced by a sample of nucleosome-free, fragmented genomic DNA. For immunoprecipitation experiments, it is additionally important to account for non-specific binding during sample preparation to obtain meaningful correlation functions (**Fig. S2** *B*). This is of increasing importance for decreasing signal-to-background ratio (**Fig. S2** *C*). The appropriate control *C* can be obtained from an immunoprecipitation with a non-specifically binding antibody (e.g. IgG control) or from a sample that lacks the antigen of interest (e.g. a knockout cell line). We devised the following strategy to compute normalized occupancy profiles that were used in the subsequent analysis. First, the normalized coverage of the control *C*_norm_ and of the specific immunoprecipitation *A*_norm_ were obtained by dividing by input signal *I* according to Eq. 1:

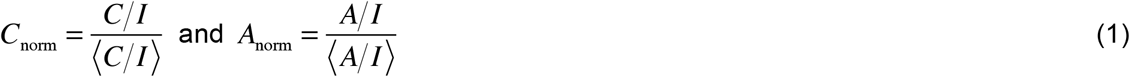

Here, (…) denotes averaging along the genomic coordinate. For the calculation of coverage (*C /1* and *A /1*) and average values ((*C/I*) and (*A/I*)), positions with zero input coverage were neglected. Subsequently, the coverage at these positions was set to the respective average value ((*C/1*) or (*A/I*)) that was calculated for the remaining positions, thus eliminating fluctuations and corresponding contributions to the correlation coefficient from these positions. In the next step, non-specific background signal was removed to obtain the normalized read occupancy *O*:

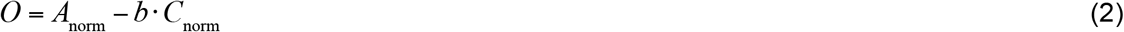

In Eq. 2, the parameter *b* quantifies the contribution of the control signal present as background in the sample (IP). To estimate *b*, we minimized the absolute value of the Pearson correlation coefficient *r*_0_ at zero shift distance between the normalized occupancy *O*and the control coverage *C*_norm_ according to Eq. 3:

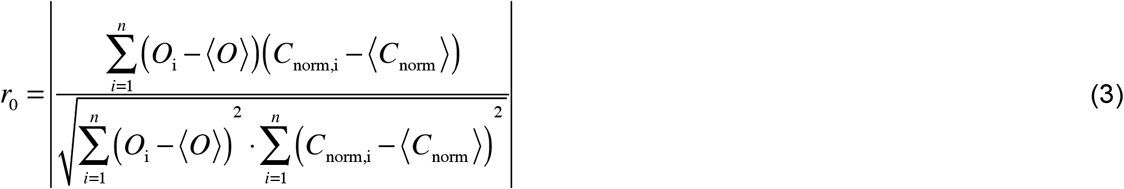

Here,*n* denotes the maximum genomic position considered for the calculation, which is typically the chromosome length. For the minimization procedure, *b* was changed between 0 and 1. Because the minimum correlation *r*_0_(*b*) indicates the lowest similarity between normalized occupancy profile and control, the corresponding *b* value was used for normalization according to Eq. 2.

### Computation of correlation functions

The Pearson correlation coefficient *r* at shift distance Δχ was calculated for the corrected data sets after shifting the two occupancy profiles *O*_1_ and *O*_2_ with respect to each other by Δχ base pairs according to Eq. 4 (similar to Eq. 3 but with a second shifted occupancy instead of the control coverage):

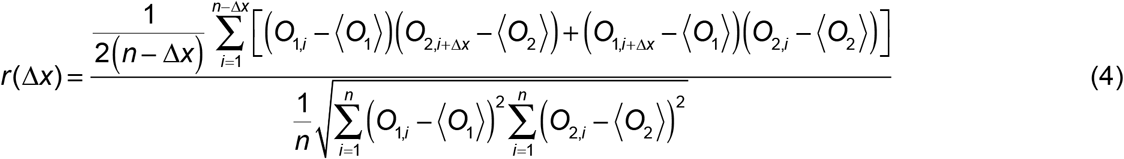

To sample the correlation function in a quasi-logarithmic manner (33), profiles were binned by a factor of 2 after 25 shift operations to double the step size. To preserve high resolution for small shift distances, the first binning operation was carried out at a shift of *Δχ* = 50 bp. This calculation was done for each chromosome separately because continuous domains cannot exceed chromosomal ends. Most correlation functions shown in the manuscript refer to chromosome 1, which is representative for all chromosomes as judged by the relatively small deviations among chromosomes (**Figs.2**,*A* and *B*, and **S8** *B*). However, correlation functions can also be calculated for smaller genomic regions (see **Fig. S1** for the correlation function for a single domain).

To compare cross-correlation functions between different features, normalization to the geometric mean of the two replicate correlation functions was conducted according to:

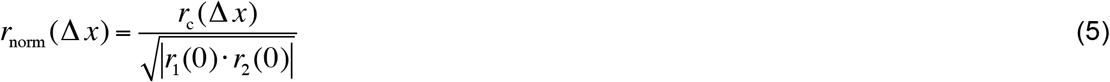

Here, *r*_c_ is the cross-correlation coefficient at a given shift distance Δχ, and *r*_1_ and *r*_2_ are the replicate correlation coefficients of the data sets used. This normalization step accounts for differences in the distributions of the features involved. For calculating the cross-correlation functions between two different features or the same feature in two different cell types at least two replicates for each sample were used. Accordingly, a cross-correlation function for each combination of replicates was computed, which results in *n^2^* functions for *n* replicates of each sample, and the average of these correlation functions was reported.

### Statistical analysis of correlation functions

Statistical analysis of data was conducted by computing standard errors and 95% confidence intervals. To assess significance and associated errors/confidence intervals for a given correlation function we considered several types of errors:
*Statistical error of the computed correlation function*. Because correlation functions are calculated from millions of regions they typically have a very small statistical error. The sample size *N* for each shift distance Δχ is given by the distance between the first and last position that is covered on the chromosome (*P*_min_ and *P*_max_) subtracted by the shift length (Δχ) according to *N(Ax)* = *P*_max_-*P*_min_-Δχ. Based on the sample size, 95% confidence intervals can be obtained using the Fisher transformation (**Fig. S8** *A*) (34, 35). If normalized occupancy values *O*_i_ follow a normal distribution reasonably well (**Fig. S8** *D*), the Fisher transformation is a good way to rapidly estimate confidence intervals for correlation coefficients. An alternate non-parametric option that is compatible with arbitrary sample distributions is bootstrapping (36). In this case, occupancy profiles are resampled with replacement in pairs (*O*_1,i_, *O*_2_,_i+Δχ_) and subsequently used for calculation of the correlation coefficient according to Eq. 4. This procedure is repeated multiple times to obtain a distribution of correlation coefficients for every pair of resampled occupancy profiles (**Fig. S8** *E*) and every shift distance Δ*χ*. Based on the width of this distribution estimates for confidence intervals are obtained. For the cases tested here, bootstrapping yielded moderately larger confidence intervals than those obtained using Fisher transformation, but intervals from both methods were of the same order of magnitude (**Fig. S8** *F*).

*Variation among chromosomes*. An estimate for the error of genome-wide domain structures or positional relationships can be obtained by comparing correlation functions calculated for different chromosomes as shown in **Fig. S8** *B*. If the relationship is governed by the same biological mechanism on all chromosomes this variation can be used to evaluate the error.

*Reproducibility of experiments*. Sample preparation might introduce a global bias into a given data set. This is generally true for deep sequencing experiments irrespectively of which method is used for downstream analysis. Such variations among replicates might not be captured by statistical comparisons conducted on the basis of a single data set or a pair of data sets. Experimental reproducibility can be assessed with MCORE for data sets with at least three different replicates by computing the correlation function for all possible combinations of samples, i.e. *n·(n-1)/2* correlation functions for *n* replicates. Subsequently, average and standard errors are calculated. We found this approach to be particularly useful to identify variations due to different experimental conditions. For example, we evaluated the changes of ChIP-seq results after using antibodies from different companies (**Fig. S10**).

*Statistical comparison of two correlation functions*. After correlation functions, associated errors and confidence intervals have been computed two functions can be compared according to standard statistical tests. An *R*-script that uses a t-test to assess the difference between two functions for each shift distance Δχ (**Fig. S8** *G*) is included in the Supporting Material.

### Quantification of MCORE correlation functions

Correlation functions obtained by MCORE provide information on the overall degree of (anti-)correlation between two deep sequencing data sets but also reflect the underlying chromatin domain structure with respect to (i) the number of chromatin domains, (ii) the relative domain abundance, (iii) the length of the respective domains, and (iv) the nucleosome repeat length. To extract the domain size distribution of a given chromatin feature, two different strategies were implemented in MCORE, which differ in the level of complexity but yield similar information. The first approach is independent of user-defined settings and computes parameters for the domain size distribution from the inflection points of the correlation function in logarithmic representation and a Gardner transformation of the correlation function. The Gardner transformation characterizes the decay spectrum of a function in a non-parametric manner (37). This workflow represents a robust approach to evaluate genome-wide features from deep sequencing data without input parameters. In particular, inflection points are completely model-independent, whereas the Gardner spectrum makes the generic assumption that the decay spectrum can be approximated by a superposition of exponential functions. The second approach can be used to quantitatively describe the domain size distribution based on a fit function. For this purpose, it is crucial to avoid over-fitting of the data. Accordingly, we implemented a complementary set of four fit options that allow for an in-depth analysis of correlation functions reporting fit parameters and their errors, thus determining domain sizes and their relative abundance. The performance of the different fit approaches is described below and in the MCORE software manual. The workflow we used in this manuscript is validated with simulated data in **Fig. S7**. *Least-squares spectrum fit*. The exponential decay spectrum for the correlation function is optimized by conventional non-linear least squares fitting. The amplitudes for a given number of (logarithmically spaced) domains are optimized to obtain a good fit. The goal of the spectrum fitting process is it to determine the length scales that are present in the decay spectrum of the curve. To this end it is not always necessary to exactly describe the shape of the correlation function. For example, the initial decay of the function is frequently too steep to be adequately fitted with a superposition of exponential functions. Nevertheless, decay lengths are typically obtained in a reliable manner. The multi-exponential fit described below often performs equally well in identifying length scales and provides a good description of the correlation function. Thus, the least-squares spectrum fit is only recommended if the multi-exponential fit does not converge properly, i.e., if it yields length scales that are very different from those determined by inflection points.

*Maximum entropy method (MEM) spectrum fit*. The exponential decay spectrum is fitted similar to the least-squares method. However, the entropy of the amplitude spectrum is maximized along with the fit quality. To this end, optimization is carried out in a parameter space that is spanned by the first derivative of the entropy and the first and second derivatives of the fit quality according to the approach described previously (38). This fit option is only recommended if the number of components obtained from the least-squares spectrum fit is much larger than the number of inflection points.

*Multi-exponential fit implemented in MCORE*. For multi-exponential fitting the following equation consisting of a combination of exponential functions is used:

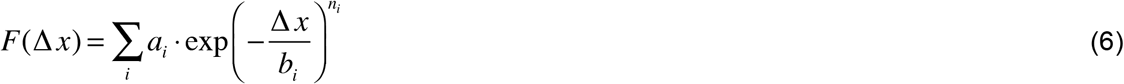

The exponential terms describe the domain structure of the correlation function, with *a*_i_, *b*_i_ and *n*_i_ yielding the relative abundance, the half width and the fuzziness of the *i*-th domain, respectively. Small exponents *n*_i_ correspond to long-tail decays in the domain size distribution.

*Multi-exponential fit in R*. The multi-exponential fit implemented in *R* (39) uses a sum of exponential functions (see Eq. 6) multiplied with an additional oscillatory term to describe the correlation function:

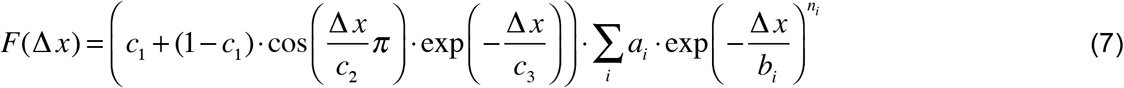

The oscillatory term accounts for the nucleosomal pattern, with parameters *c*_1_ for the strength of the nucleosomal oscillation, *c*_2_ representing the nucleosomal repeat length and *c*_3_ the scale on which regular nucleosomal spacing is lost. When using this approach, the minimal number of exponential terms that yielded uncorrelated fit residuals was chosen.

### MCORE runtime

Generation of normalized occupancy profiles and calculation of the respective correlation function for the entire chromosome 1 takes 15–20 minutes on a laptop computer with a 2.7 GHz Intel Core i5 processor and 8 GB memory. For smaller chromosomes or genomic regions of interest the calculation is faster.

### Peak calling

Peak calling was done using MACS (10) and SICER (11). Prior to peak calling reads were preprocessed as described above including mapping to the mouse mm9 assembly by Bowtie (32), considering only uniquely mapping hits without mismatches and removing duplicates. Peak calling was done using default parameters and the input as control file. For H3K36me3 MACS mfold levels 5, 10 and 30 were tested, and mfold 5 was selected. For SICER the FDR threshold was set to 0.0001, a window size of 200 bp and a gap size of 600 bp were used for H3K9me3 and H3K36me3, and a window size of 200 bp and a gap size of 200 bp were used for H3K4me3.

### Network models

Graphs for network models were created and plotted using Gephi (http://gephi.github.io). Nodes were manually prearranged, and their layout was optimized using the Fruchterman-Reingold algorithm (40), which adjusts node positions based on forces that act between nodes according to the respective correlation strength.

### Sample preparation for histone ChIP-seq

ESCs and neural progenitor cells from 129P2/Ola mice were cultured and differentiated as published (41). ChIP-seq experiments and mapping of reads to the mm9 assembly of the mouse genome was conducted as described previously (22). In brief, 10^6^ cells were cross-linked with 1% PFA and cell nuclei were prepared. Chromatin was sheared by sonication to mononucleosomal fragments. ChIP was carried out with antibodies (Abcam) against H3K4me1 (ab8895), H3K4me3 (ab8580), H3K9me3 (ab8898), H3K27ac (ab4729), H3K27me3 (ab6002), H3K36me3 (ab9050) or an unspecific IgG from Acris (RA073 or PP500P) (**Table S5**). Libraries were prepared according to Illumina standard protocols with external barcodes and were sequenced with 51 bp single-end reads on an Illumina HiSeq 2000 system. After sequencing, cluster imaging and base calling were conducted with the Illumina pipeline (Illumina). 20 - 30 Mio reads were obtained for each sample. Reads were uniquely mapped without mismatches to the mm9 mouse genome using Bowtie. For RNA-seq, cells were harvested and long RNAs were isolated with the RNeasy Mini Kit (Qiagen), DNA was digested by DNase I (Promega) for 30 min at 37°C, and libraries were prepared using the Encore Complete RNA-Seq Library Systems (NuGEN).

### Data and software

ChIP-seq data have been deposited to the GEO database under the accession number GSE61874. An executable Java program, including a test data set and an *R* script for statistical testing of the difference between two correlation functions, is available in the supplemental material and can be downloaded at http://malone.bioquant.uni-heidelberg.de/software/mcore.

## Results

### Comparison of MCORE to other sequencing analysis workflows

The MCORE workflow in comparison to the currently most common approaches for deep sequencing analysis is illustrated in **Figs. 1** and **S2** *A.* First, all types of data sets were transformed into normalized read occupancy profiles. Among others, this normalization step takes into account the propensity of a DNA fragment to be ligated, amplified, sequenced and mapped. To correct for these multiplicative biases, the sample read density was divided by the input read density for immunoprecipitation (IP) and Hi-C experiments or by the sum of converted and unconverted read densities for bisulfite sequencing (BS-seq). We expect that Hi-C data that have already been normalized with other methods (42, 43) in a similar manner can be used for MCORE without further correction. IP experiments such as ChIP-seq yielded significant background correlation due to non-specific binding of DNA and proteins to beads or bead-antibody complexes (44). Accordingly, these data sets were further corrected by subtraction of a weighted control IP signal obtained from an IP with non-specific antibodies (**Fig. S2** *B*). The weighting factor reflects the contribution of non-specifically precipitated DNA in each sample and removes the correlation between specific IP and control IP (Materials and Methods). As expected, the contribution of non-specific signal depended on the quality of the antibody and on the enrichment levels of the specific IP-signal. H3K9me3 ChIP-seq data, for example, were affected more strongly by this correction than H3K4me3 ChIP-seq data (**Fig. S2** *C*), because H3K4me3 domains were more distinct and exhibited larger enrichment levels than H3K9me3 domains. Normalized occupancy profiles can be exported and also be used for other downstream analysis methods.

Peak calling or dynamic network models use occupancy profiles from mapped reads to define peaks or chromatin states based on local enrichments (**Fig. 1** *A*). In contrast, MCORE computes correlation functions from the sequencing read occupancy without binarizing the data. To this end, normalized occupancy profiles from two different data sets were shifted with respect to each other along the genomic coordinate, and the normalized Pearson correlation coefficient for each shifting distance Δχ was calculated and analyzed (Materials and Methods). In contrast to rank correlations the Pearson correlation coefficient accounts for the enrichment values within the normalized occupancy profile and therefore preserves the biologically relevant information (**Fig. S3**). We computed three types of correlation functions with different biological meaning: (i) the correlation function between two replicates, yielding the domain topology for a chromatin feature (**Fig. 1** *B*), (ii) the correlation function between the same feature in two different cell types, providing information on the positional conservation of a given chromatin mark across cell types (**Fig. 1** *C*), and (iii) the correlation function between two different features in the same cell type, reflecting their genome-wide positional relationship such as co-localization or shifted localization (**Fig. 1** *C*). The use of at least two independent data sets (either two replicates or two samples interrogating different features or cell types, see Eq. 4) for the calculation of each type of correlation function suppresses spurious noise that is uncorrelated between independent experiments and does therefore not contribute to the correlation.

**Figure 1.**
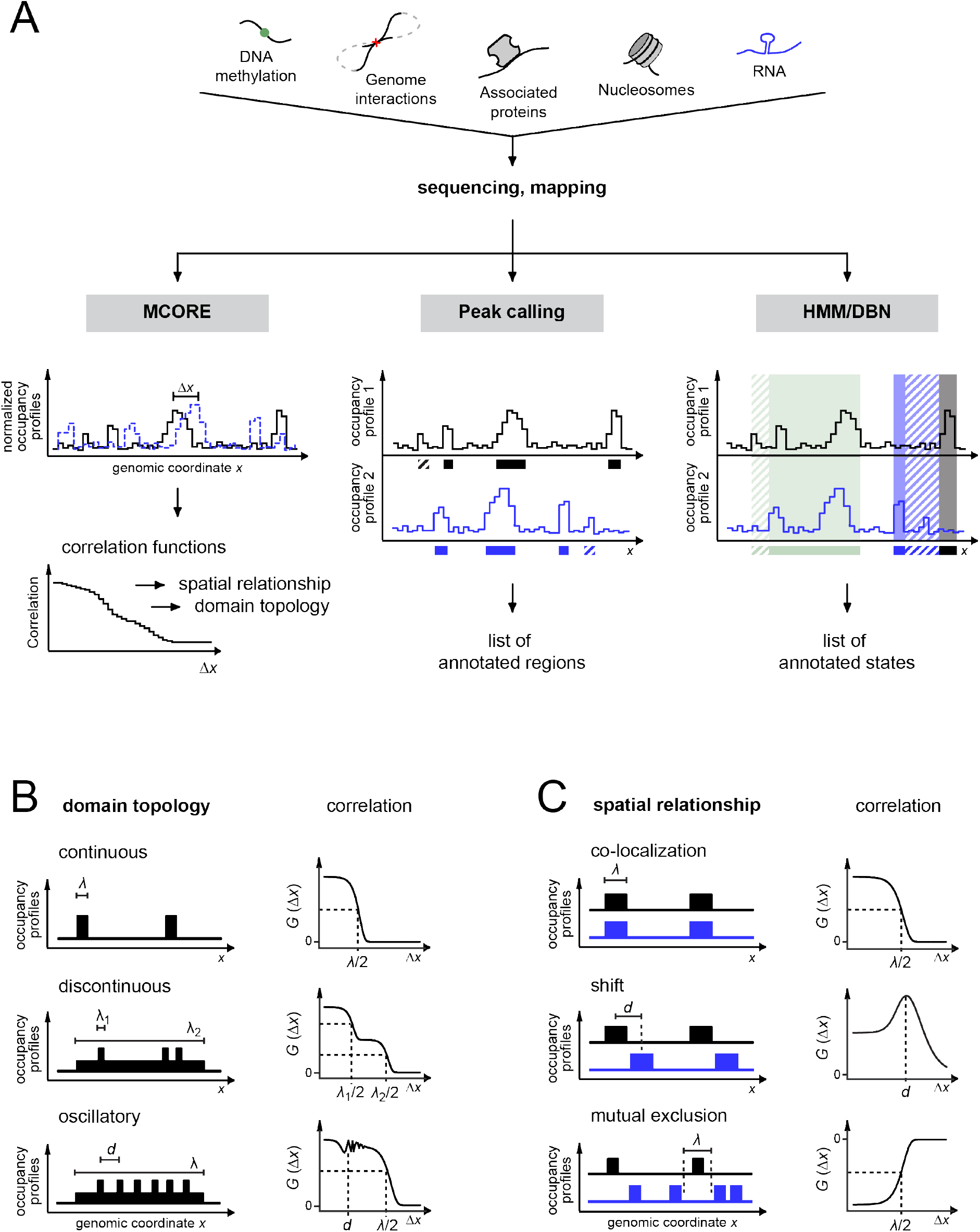
MCORE can identify and compare patterns in deep sequencing data sets. **(A)** MCORE is suited for the analysis of deep sequencing data from various methods. Initially, mapped reads are used to compute occupancy profiles of two samples (black/blue). Subsequently, in case of MCORE the profiles are normalized using the input sample and, if applicable, the control sample. In contrast to other methods like peak calling, hidden Markov models (HMM) or dynamic Bayesian networks (DBN), which use control and IP samples for the detection of enriched regions, MCORE does not score enriched regions but rather shifts normalized occupancy profiles with respect to each other to compute correlation functions, which contain information about chromatin patterns as illustrated in panels *B* and *C* and **Fig. S1.** To this end it uses all sequencing reads without filtering and avoids any assumptions about the enrichment pattern. **(B)** Correlation functions between replicates for the same chromatin feature contain information about its domain topology. Whereas the correlation coefficient at shift distance zero quantifies the reproducibility of the experiment, the shape of the function reflects the distribution of the feature along the genomic coordinate. Continuous domains lead to a steep decay at the shift distance that coincides with half the domain size *λ* (top), whereas broad domains containing small highly enriched regions yield multiple decay lengths *λ*_i_ (center). Arrays of equally spaced domains cause an oscillating contribution in the correlation function (bottom). Mixtures of domains with different topology yield a superposition of the respective correlation functions. **(C)** Correlation functions between two different chromatin features reflect their spatial relationship. Co-localizing features yield monotonously decaying functions (top) that resemble those between replicates discussed in the previous panel. Correlation functions for features that are shifted with respect to each other exhibit a local maximum at the shift distance *d* (center). Mutually exclusive features are recognized by negative correlation amplitudes (bottom). Features that do not exhibit any spatial relationship with respect to each other yield no correlation for any shift distance.

To compare co-localization values among differently distributed marks we normalized cross-correlation functions with respect to their replicate correlation (Materials and Methods, Eq. 5). This step was required because broadly distributed marks tended to yield smaller cross- and replicate correlation coefficients than marks forming narrow and well-positioned domains. As illustrated in **Fig. 1** *C*, positive correlation indicated co-localization at a given shift distance, whereas negative correlation reflected mutually exclusive modification or binding. Each decay length and its contribution to the correlation function encoded a domain size and its abundance, whereas superimposed oscillations reflected nucleosome spacing (31, 41). Where necessary, the correlation function can be used as a starting point to identify individual regions of interest as described below.

MCORE is complementary to peak calling, which generally aims to identify enriched regions without larger gaps. As the probability to find modified regions without spurious gaps decreases with size, broad regions are prone to get lost or fragmented in such analyses. This phenomenon is more or less pronounced depending on the settings and the algorithm used as shown for H3K9me3 in **Fig. S4** *B*. Further, it is often challenging to identify and remove false-positive/negative peaks that are caused by the inherent properties of sequencing data sets like noise, artificial overrepresentation of particular genomic regions (45, 46) or insufficient read coverage (15). An example for H3K36me3 is shown in **Fig. S4** *C*. MCORE retrieves information about patterns upstream of peak calling analyses and is relatively robust towards uncertainties at individual loci because correlation functions are calculated from the entire collection of sequencing reads in a large genomic region (see **Figs. S5** and **S6** for the influence of read coverage).

### Interpretation and quantification of correlation functions

We quantified the information contained in correlation functions by first analyzing their decay spectrum in a model-independent manner and by subsequently fitting a generic model function (29) as described in the Materials and Methods section. This is illustrated for a simulated data set in **Fig. S7**. As a first step, inflection points (in logarithmic representation) were numerically determined, yielding the decay lengths that are present in the correlation function. Depending on the type of function these decay lengths *λ*_i_ represent domain sizes or separation distances (**Fig. 1** *C*). Next, the Gardner transformation was computed, which exhibited peaks at the characteristic decay lengths (37). Both approaches were independent of input parameters or model assumptions. Finally, we fitted the correlation function to quantitatively describe the domain size spectrum (Materials and Methods). Because decay lengths and nucleosome repeat length follow from the change of the correlation coefficient with shift distance, these parameters are independent of the absolute correlation amplitude, which is beneficial for the analysis of data sets that are not properly normalized, e.g. due to low sequencing depth or lack of suitable control samples.

Correlation functions can be compared to each other based on errors obtained from Fisher transformation or bootstrapping (**Fig. S8**, Materials and Methods). These errors reflect variations of the correlation coefficient among different positions within the genomic region of interest. If more than two replicates were available, replicate correlation functions calculated for each combination of independent samples were combined to account for differences among experiments (**Fig. S8**). We found these errors most meaningful because the variability among replicates can typically not be neglected and should be used as a reference when comparing different correlation functions to each other. The shape and the amplitudes of correlation functions were well reproducible when normalized according to the workflow described above. This was also true when comparing our samples with published histone modification ChIP-seq samples from other labs (**Figs. S8** *C* and **S9** *A*).

In summary, MCORE yields compact genome-wide representations of chromatin features in the form of correlation functions that can be quantitatively evaluated and compared to each other. It can be used to (i) determine domain topologies (**Fig. 1** *B*), (ii) assess positional relationships (**Fig. 1** *C*), (iii) test the reproducibility of experiments, or (iv) assess variations caused by changes in experimental conditions, e.g. the use of antibodies from different suppliers (**Fig. S10**). In contrast to the Pearson correlation coefficient between two data sets alone, the normalized correlation function provides insight into the similarity of the data sets on a broad range of length scales. Thus, MCORE can detect changes in domain size, amplitude or relative genomic position and can be used to track the reorganization of the epigenome among different cell types as shown below.

### Domain structure and nucleosome pattern of modified regions in ESCs and NCs

We used replicate correlation functions to dissect the domain structures and nucleosome patterns in ESCs and NCs throughout the genome (**Figs. 2**, *A* and **B**, and **S11; Tables S3** and **S4**). These quantities reflect the activity of the cellular machinery that shapes the chromatin landscape and thereby regulates chromatin function. Most features studied here, such as H3K9me3, displayed complex domain size distributions with multiple characteristic decay lengths (**Fig. 2**, *A* and *B*). An exception was H3K4me3, which in agreement with published data (47) formed almost exclusively distinct peaks of roughly 1900 bp or 9–10 nucleosomes in size in both ESCs and NCs. For H3K36me3 we found a typical domain size of 24–30 kb, which is of the same order of magnitude as the average gene length in the mouse genome (according to NCBI Build 37, mm9). The nucleosome repeat length varied among domains carrying different histone modifications, with 218 bp for H3K27me3 in NCs and 182 bp for H3K9me3 and H3K36me3 in NCs (**Tables S3** and **S4**). This observation suggests that nucleosome spacing is differentially regulated and linked to the chromatin state, consistent with previous reports (31, 48).

**Figure 2.**
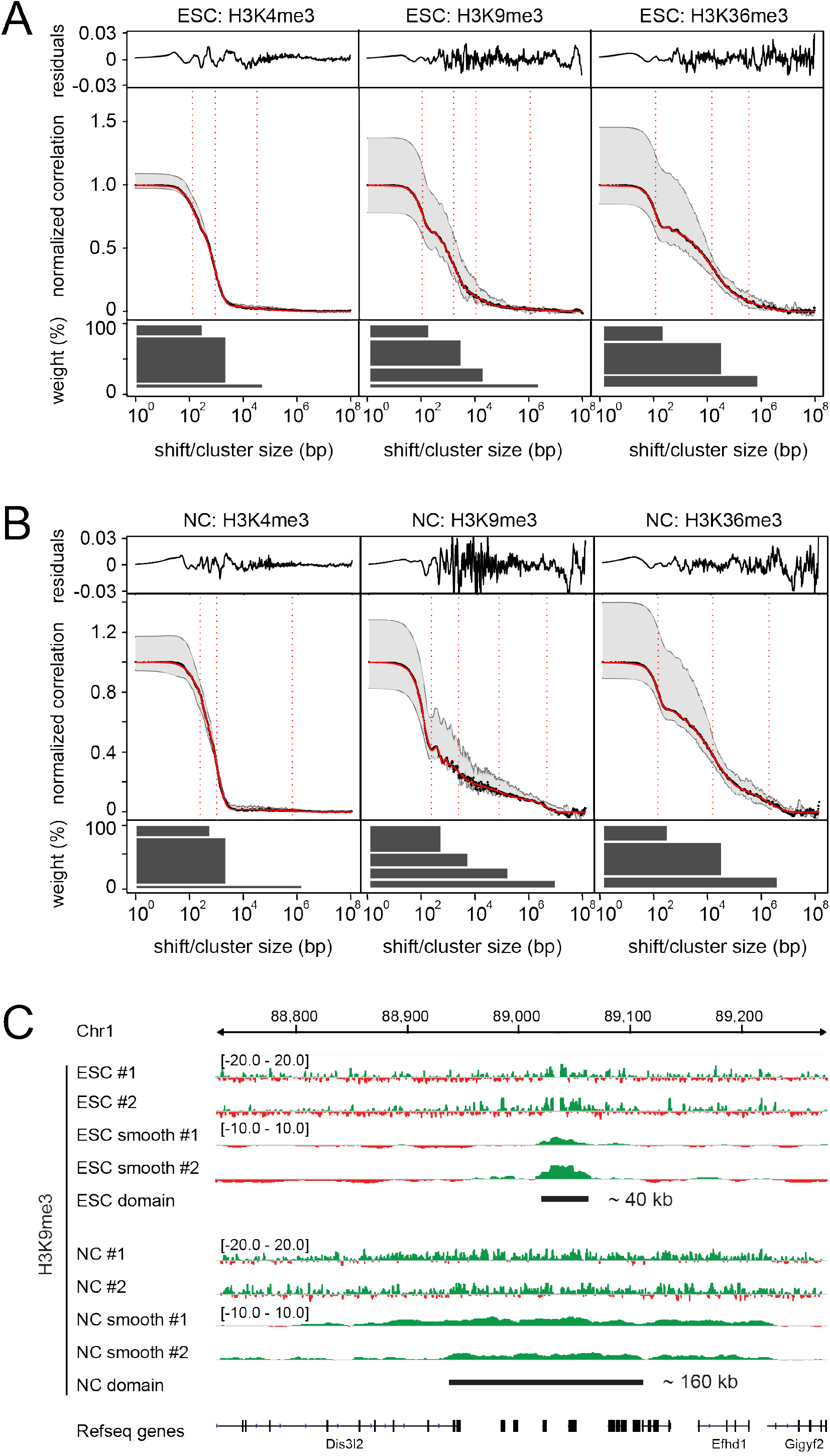
Quantification of domain sizes for different histone marks. **(A)** Correlation functions for replicates in ESCs. Correlation functions calculated between replicates for chromosome 1 (black) and their fit functions (red) with characteristic domain sizes obtained from the fit (vertical dotted lines) are shown. Gray regions indicate maximum variation between chromosomes. Fit residuals are plotted above the correlation curves. Domain sizes and abundances calculated from the respective fit parameters are shown below the correlation curves. **(B)** Same as in panel *A* for NCs. **(C)** As shown in panels *A* and *B*, MCORE identified broad H3K9me3 domains spanning on average 128 kb and 7.6 Mb in NCs, which were absent in ESCs. To annotate the genomic positions of these domains, read counts in a sliding window of 128 kb, which corresponded to the smaller domain size, were evaluated. An example of a domain that became broader in NCs is shown (‘#1’ and ‘#2’ denote replicates). For clarity, the occupancy profiles were smoothed with 0.2-times the window size (‘smooth’). For window size 7.6 Mb see **Fig. S13**.

The initial decay of most replicate correlation functions is caused by the reduced probability to find the same modification at the neighboring nucleosome and is therefore associated with a domain size of a single nucleosome. Notably, a prerequisite for this interpretation is that the occupancy profile is properly normalized and not heavily undersampled, which is validated for representative profiles in **Figs. S5** and **S6**. Accordingly, homogenous domains that primarily contain equally modified nucleosomes produce a weaker initial decay than domains that contain a mixture of modified and non-modified or differently modified nucleosomes. Whereas the subtle initial decay for H3K4me3 in ESCs and NCs (**Fig. 2**, *A* and *B*; **Tables S3** and **S4**) is indicative of homogenous domains, the pronounced decay for H3K9me3 in NCs (**Fig. 2** *B* and **Table S4**) suggests that this modification forms discontinuous domains with gaps. This is corroborated by the absence of isolated nucleosomes with high H3K9me3 enrichment levels outside broader domains (**Fig. S12**), which could also be responsible for a steep decay in the correlation function because such nucleosomes would have unmethylated neighbors.

In summary, these results indicate that different histone modifications form domains with different size and structure. Based on the domain size and frequency distribution obtained from MCORE, an assignment to specific genomic loci can be made, e.g. by evaluating the normalized occupancy profiles with a sliding window corresponding to a domain size of interest. This procedure is illustrated in **Figs. 2** *C* and **S13** for broad H3K9me3 domains, which according to MCORE prevailed in NCs.

### Changes in chromatin patterns during stem cell differentiation

To identify changes of chromatin features during stem cell differentiation we conducted a comparative MCORE analysis of more than 60 deep sequencing data sets from ChIP-seq (histone modifications: H3K4me1, H3K4me3, H3K9me3, H3K27ac, H3K27me3, H3K36me3, binding sites of RNA polymerase II (RNAP II) and transcription factors TAF3, Oct4 and Otx2), BS-seq, RNA-sequencing (RNA-seq), Hi-C and RNAP II ChIA-PET experiments in ESCs and NCs (**Figs. 2, 3, S14–17; Tables S2** and **S5**). Normalized correlation amplitudes at zero shift distance were assembled into a matrix (**Fig. 3** *A*, red/blue), reflecting co-localization or mutually exclusive localization of different features. In both cell types we found more co-localizations than mutual exclusions, which suggests that the set of chromatin features analyzed here tends to localize to the same part of the genome. In general, mutual exclusions were weaker than co-localizations as judged by the absolute values of the respective normalized correlation coefficients.

**Figure 3.**
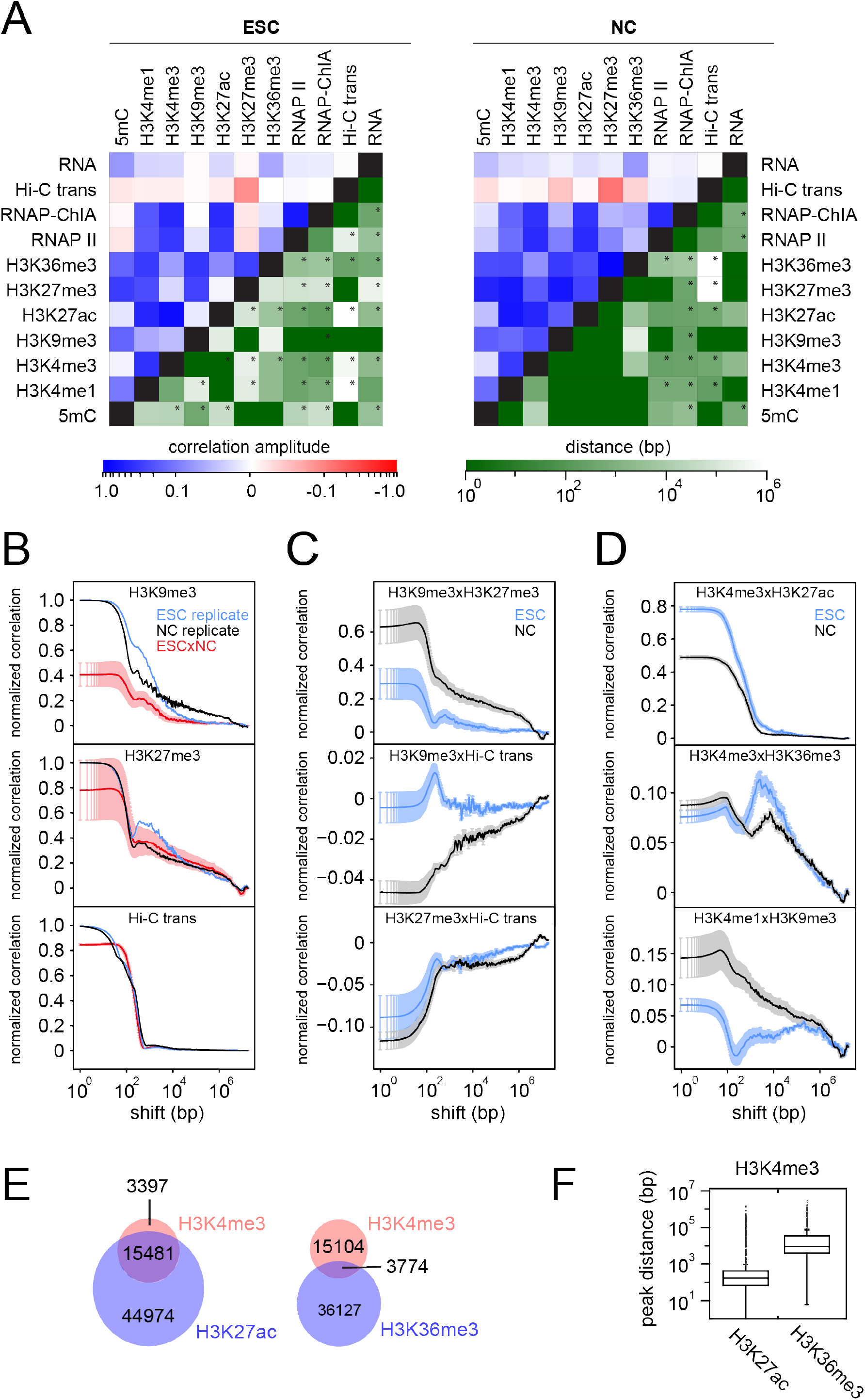
MCORE reveals genome-wide relationships between chromatin features. **(A)** Co-localization (top, red/blue) and separation distance (shift distance for the largest local maximum, bottom, green) between pairs of different features in ESCs (left) and NCs (right) are shown. Stars indicate correlation functions for which the local maximum is also the global maximum. Hi-C trans, Hi-C inter-chromosomal contacts; RNA, RNA-seq; RNAP II-ChIA, RNAP II ChIA-PET. **(B)** Correlation functions for replicates of H3K9me3, H3K27me3 and inter-chromosomal contacts (Hi-C trans) in ESCs (blue) and NCs (black) show the spatial extension of these features. Average cross-correlation functions (red) between ESCs and NCs quantify the co-localization of a given feature across cell types. Averages were calculated from the four possible combinations of the two replicates for each sample (Materials and Methods). Error bars, s.e.m. **(C)** Cross-correlations between H3K9me3 and H3K27me3 (top) or H3K9me3/H3K27me3 and inter-chromosomal contact sites (Hi-C trans, center/bottom) in ESCs and NCs. Repressive domains co-localize in NCs (top) and have a tendency to be depleted for inter-chromosomal contacts (bottom). Error bars, s.e.m. **(D)** Cross-correlations between H3K4me3 and H3K27ac (top) indicate co-localization of both marks in small domains, whereas cross-correlations of H3K4me3 and H3K36me3 (center) reveal a relative displacement of roughly 5 kb between the two marks. In NCs, there is an additional co-localization at zero shift distance that is weaker in ESCs. Cross-correlations between H3K4me1 and H3K9me3 (bottom) show that both marks are stronger co-localized in NCs than in ESCs. The local maximum at 100 kb shift distance in ESCs suggests a separation of H3K4me1 from broad H3K9me3 domains. Error bars, s.e.m. **(E)** Peak calling in NCs as readout for co-localization. Red, peaks called by MACS for H3K4me3; blue, peaks called by SICER for H3K36me3 or by MACS for H3K27ac. The numbers of (overlapping) peaks are indicated. **(F)** Distribution of distances between called peaks. Distances were calculated from the center of the H3K4me3 peak to the center of the nearest peak in the second data set (H3K27ac or H3K36me3).

In ESCs, the strongest co-localizations were found among features related to actively transcribed genes (H3K4me1, H3K4me3, H3K27ac, H3K36me3, RNAP II, RNAP II ChIA-PET). Notably, H3K36me3, which is known to be associated with active genes, also co-localized with H3K9me3/H3K27me3, which are traditionally considered heterochromatin marks. This might reflect (i) the presence of repressed genes not devoid of H3K36me3 (49), (ii) the occurrence of H3K9me3 and H3K27me3 at active genes (47), and/or (iii) the presence of H3K36me3 domains outside of coding genes. Mutual exclusion was found between RNAP II and the repressive marks H3K27me3 and 5mC (but not H3K9me3) in ESCs. Furthermore, inter-chromosomal contact sites were depleted around H3K27me3 in ESCs, indicating that H3K27me3 domains localized preferentially inside chromosome territories.

In NCs, co-localization among features associated with active chromatin was conserved and tended to become stronger (**Fig. 3** *A*). Most activating modifications retained their domain size structures and genomic positions on a global level (**Fig. S15**). In contrast, H3K9me3 and H3K27me3 redistributed during differentiation in a way that their co-localization with each other, with 5mC and with some of the activating marks like H3K4me1 increased (**Figs. 3**,*A* and *D*, and **S16**). In particular, the following changes are noteworthy: (i) Both H3K9me3 and H3K27me3 formed broader domains in NCs compared to ESCs, which led to a stretched decay in correlation functions for NCs compared to the steeper decays in correlation functions for ESCs (**Figs. 2**, *A* and *B*, and **3 B**). (ii) The normalized correlation of H3K9me3 between ESCs and NCs decreased compared to the normalized correlation between replicates from the same cell type (**Fig. 3** *B*). The same tendency was observed for H3K27me3. These differences suggest partial re-location of H3K9me3/H3K27me3 during differentiation because otherwise correlation functions between ESCs and NCs would resemble the correlation function calculated for the replicates from the same cell type, and all curves in each panel would essentially be identical. (iii) The normalized correlation between H3K9me3 and H3K27me3 increased in NCs (**Fig. 3** *C*), which is indicative of stronger co-localization of both marks in NC ensembles. (iv) Correlation functions for 5mC in ESCs, NCs and between both cell types were similar (**Fig. S15**). Thus, global changes in the genome-wide 5mC pattern were minor, consistent with previous findings (47). (v) The normalized correlation between H3K27me3 and 5mC was higher in NCs compared to ESCs (**Figs. 3** *A* and **S17** *A*), suggesting re-localization of H3K27me3 to 5mC domains. Normalized correlation between H3K9me3 and 5mC increased for large shift distances in NCs, implying that extended H3K9me3 domains formed in the vicinity of pre-existing 5mC sites (**Fig. S17** *A*). (vi) Substantial mutual exclusion was found between H3K9me3 and inter-chromosomal contacts in NCs but not in ESCs, which suggests that H3K9me3 was re-localized to the interior part of chromosome territories (**Fig. 3** *C*). H3K27me3 resided preferentially inside chromosome territories already in ESCs and did not change its position in NCs (**Fig. 3** *C*).

### Differential relationships among chromatin features in ESCs and NCs

Next, we determined the characteristic genomic separation distance for each pair of features (**Fig. 3** *A*, green color coding). Whereas correlation functions for co-localizing features tend to decrease monotonously, correlation functions for shifted features exhibit local maxima at their characteristic separation distance (**Fig. 1** *C*). Correlation functions for features that co-localize at some regions in the genome and are shifted with respect to each other at other places exhibit an initial decay that is followed by local maxima (**Fig. 3**, *C* and *D*). This type of information is lost in evaluation schemes that exclusively assess overlap (**Fig. 3** *E*). For simple cases, such as H3K4me3 and H3K36me3 that localize side by side at promoters and bodies of active genes (**Fig. S4**), similar information is obtained by determining distances between adjacent peaks across data sets (compare **Fig. 3**, *D* and *F*).

Examples for pairs of features that are shifted with respect to each other in ESCs but overlap and co-localize in NCs are H3K4me1-H3K9me3 (**Fig. 3**, *A* and *D*), H3K4me3-H3K27me3 and H3K9me3-H3K27ac (**Fig. 3** *A*). These changes are in agreement with the global reorganization of H3K9me3 and H3K27me3 in NCs described above.

### Network models for relationships among chromatin features on multiple scales

The cross-correlation functions introduced above represent the scale-dependent relationships between pairs of chromatin features. Accordingly, we used these values to construct network models that reflect the associations among all features assessed here for a particular genomic distance (**Fig. 4**). Features were arranged based on their associations at zero shift distance, with positively correlated features positioned close to each other (Materials and Methods). As described above, activating histone modifications such as H3K4me1, H3K4me3 and H3K27ac co-localized with RNAP II and RNAP II ChIA-PET sites in both ESCs and NCs. Repressive marks including H3K9me3, H3K27me3 and 5mC were also positively associated with each other, with stronger correlations in NCs than in ESCs. This observation suggests that in NCs a larger fraction of the genome is heterochromatic. H3K36me3 exhibited positive correlations with both activating and repressive marks, indicating partial overlap of the respective domains. Associations among different features changed in a characteristic manner with genomic distance, reflecting the mechanisms that establish chromatin patterns on different scales. Activating features remained associated with the adjacent nucleosome (200 bp shift), indicative of chromatin domains that extend beyond a single nucleosome. In contrast, the cross-correlation among repressive marks at neighboring nucleosomes decreased considerably compared to their correlation at the same nucleosome. This points to the presence of nucleosomes (without an equally modified neighbor) that either carry at least two repressive marks simultaneously, transition between two different repressive marks over time, or stably carry different repressive marks in different cells. All of these scenarios would produce positive correlation in the ensemble average. At a shift distance of about ten nucleosomes (2000 bp), most associations among activating histone modifications were lost, reflecting the relatively limited spatial extension of the respective domains (**Tables S3** and **S4**). In contrast, correlations between repressive marks decreased only moderately, which is consistent with their occurrence in broad domains with low enrichment levels. The differential scale-dependence found for relationships among active and among repressive marks suggests distinct topologies of the respective chromatin domains and thus points to fundamental differences in the mechanisms for their establishment and maintenance.

**Figure 4.**
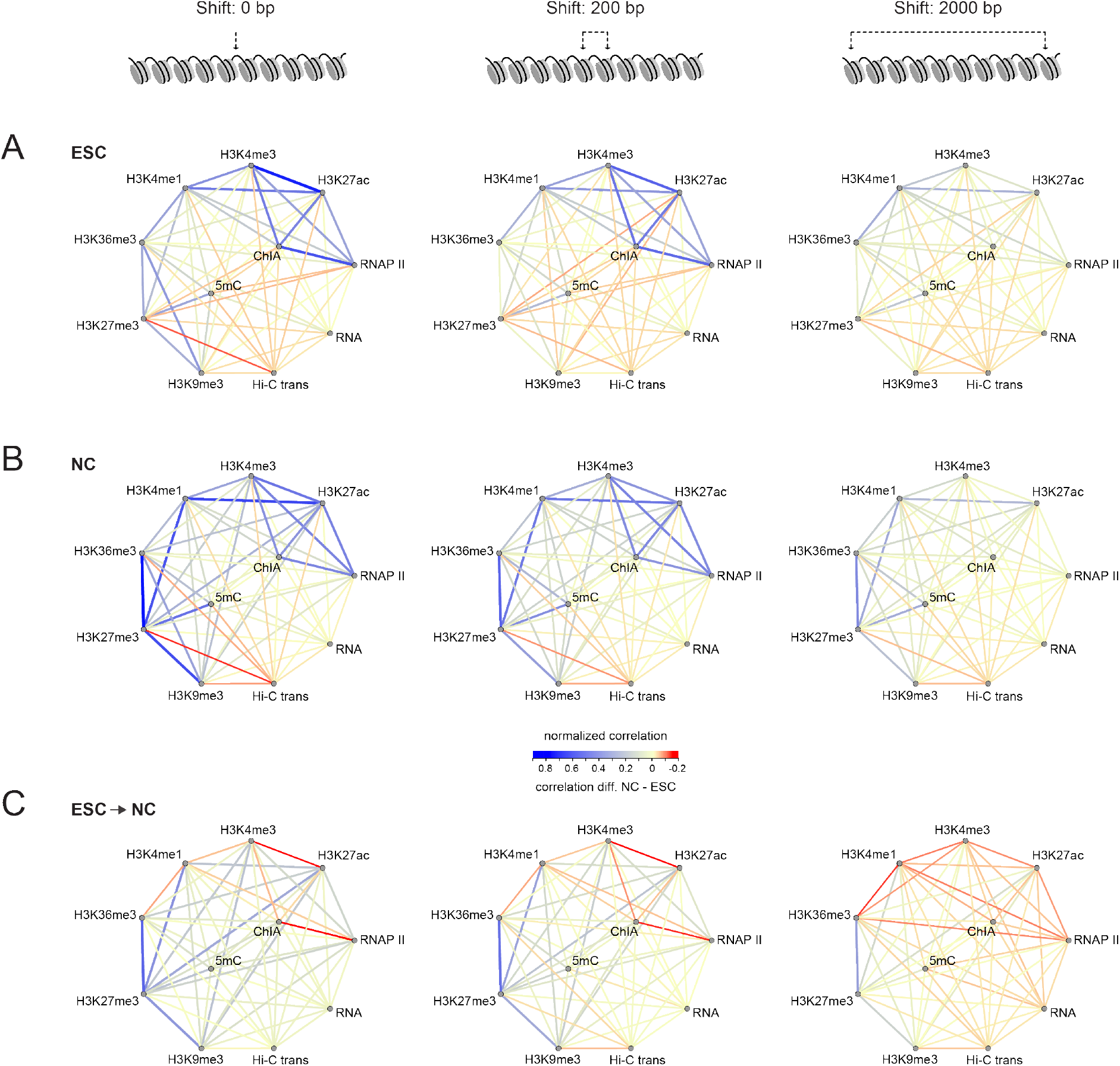
Network models for scale-dependent relationships among chromatin features. **(A)** Network models illustrating the relationships among different chromatin features in ESCs on different scales (blue: positive correlation, red: negative correlation). Features were grouped according to their correlation at zero shift distance (left), yielding a cluster of features associated with active transcription and a cluster of marks related to gene silencing, whereas H3K36me3 co-localizes with members of both groups. The correlations among features on adjacent nucleosomes (200 bp shift distance) differ from the correlations among features at the same nucleosome (0 bp shift distance), indicating that only some features form continuous domains that extend beyond a single nucleosome. For the even larger shift distance of roughly ten nucleosomes (2000 bp), only a few long-range correlations remain, which either reflect large domains of co-localizing features or features that are shifted with respect to each other. The latter two possibilities can be distinguished based on the shape of the correlation function (**Fig. 1** *C*). **(B)** Same as in panel *A* but for NCs. **(C)** Network models illustrating changing relationships among different chromatin features in ESCs and NCs. The difference NC-ESC is colored in blue if correlations became stronger in NCs and red if correlations became weaker in NCs. Positive correlations among repressive marks were stronger in NCs than in ESCs, particularly on larger scales.

### Reorganization of heterochromatin components

To further investigate the changes in heterochromatin organization during differentiation of ESCs into NCs inferred from the MCORE analysis above, we dissected the core part of the network around H3K9me3. To this end we compared the distributions of the H3K9me3 mark, the histone methyltransferase SUV39H1 that sets this mark in pericentric heterochromatin, and the heterochromatin protein 1 isoforms HP1x03B1; and HP1ß to each other. Both SUV39H1 and HP1 contain chromodomains that recognize H3K9me3, but the contribution of these interactions to their genome-wide binding profiles has not been studied comprehensively. First, we asked if the two HP1 isoforms displayed cell type-specific chromatin interaction patterns. We found that the genomic distributions of HP1a and HP1ß were different from each other in both ESCs (**Fig. 5**, *A-C*) and NCs (**Fig. 5**, *D-F*). In ESCs, HP1ß formed broader domains than HP1α (**Fig. 5** *A*) that were less correlated with H3K9me3 (**Fig. 5** *B*) but rather overlapped with H3K36me3 (**Fig. 5** *C*). This finding supports recent work, which showed that HP1ß but not HP1a is enriched in exons and essential for proper differentiation and maintenance of pluripotency in ESCs (50). The nuclear distribution of HP1ß in ESCs might be related to its function in splicing (51). In NCs, HP1α and HP1ß displayed moderate differences in their domain structure (**Fig. 5**, *D* and *G*), with a stronger preference of HP1α for broad domains. In contrast to ESCs, both isoforms strongly co-localized with H3K9me3 in NCs (**Fig. 5** *E*), in line with their well-established role as heterochromatin components in differentiated cells ((22) and references therein). Co-localization with H3K36me3 was also observed (**Fig. 5** *F*), consistent with the overlap between H3K9me3 and H3K36me3 domains in NCs found above. Next, we focused on the composition of H3K9me3 domains in NCs. Whereas H3K9me3 formed both broad and intermediately sized domains, SUV39H1 did not form intermediate domains but rather broad domains containing gaps (**Fig. 5**, *D* and *G*) as suggested by the fast decay of its replicate correlation function (**Fig. 5** *D*, red). Consistently, co-localization among HP1α/β, SUV39H1 and H3K9me3 was not found in intermediate but rather in broad domains (**Fig. 5** *E*). These findings point to the presence of SUV39H1-independent H3K9me3 domains with intermediate size in NCs, which have also been described in ESCs (52), indicating that H3K9me3 is not sufficient for stably recruiting SUV39H1 or HP1 to chromatin. This is in line with a looping model in which well-separated high-affinity binding sites (nucleation sites), which reside within broad heterochromatic regions, recruit SUV39H1 to establish and maintain H3K9me3 (**Fig. 5** *H*).

**Figure 5.**
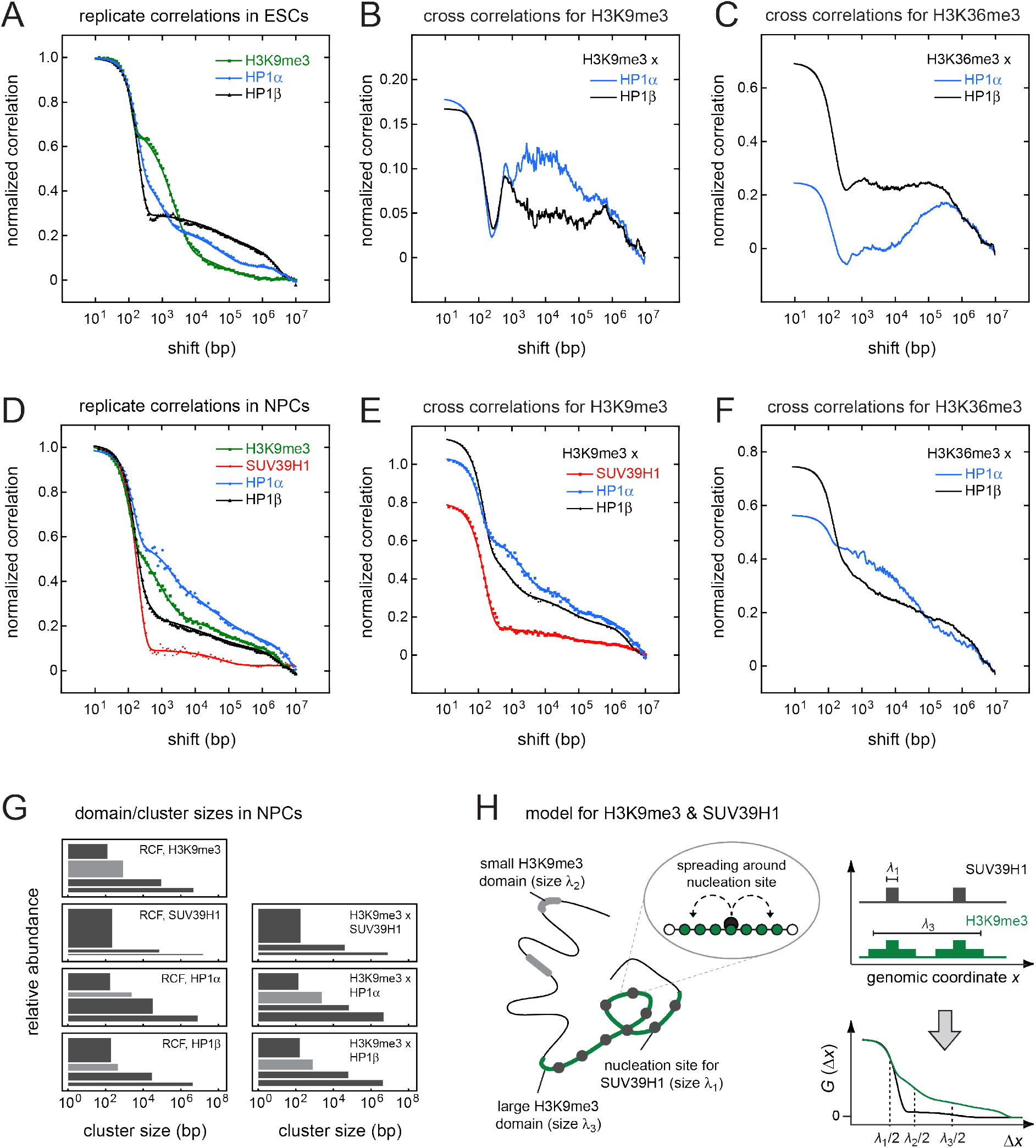
Interplay among H3K9me3, SUV39H1 and HP1. **(A)** Replicate correlation functions of HP1α (blue), HP1β (black) and H3K9me3 (green) in ESCs. **(B)** Cross correlation functions of HP1α (blue) or HP1β (black) with H3K9me3 in ESCs. **(C)** Cross correlation functions of HP1α (blue) or HP1β (black) with H3K36me3 in ESCs. **(D)** Same as in panel *A* but for NPCs and including SUV39H1 (red). H3K9me3 and HP1α/β exhibit small, intermediate and (very) broad domains. The short domain size of one nucleosome is present in the correlation functions for all marks, suggesting that domains consist of nucleation sites and gaps (as explained in the text). SUV39H1 does not form intermediately sized domains. **(E)** Same as in panel *B* but for NCs and including SUV39H1 (red). SUV39H1, HP1α, HP1ß and H3K9me3 strongly co-localized. Intermediate domains are not present in the cross correlation function between SUV39H1and H3K9me3, indicating that both features only co-localize in short and broad domains. In contrast, HP1α and HP1β essentially follow the H3K9me3 distribution, indicating that they do not distinguish between differently sized H3K9me3 domains. **(F)** Same as in panel *C* but for NCs. **(G)** Domain size distribution for correlation functions in panels *D* and *E*. **(H)** Schematic illustration of a nucleation-and-looping mechanism for the formation of SUV39H1-dependent H3K9me3 domains, which is consistent with the MCORE results for NPCs.

### Model for changes of chromatin features during differentiation

The MCORE results on domain size distributions, co-localizations and separation distances (**Figs. 2–4**) lead us to a model for the reorganization of chromatin during differentiation of ESCs into NCs depicted in **Fig. 6**. H3K9me3 and H3K27me3 domains became larger and stronger co-localized with sites of preexisting 5mC during the transition from ESCs to NCs (**Figs. 3**, *>B* and *C*, and **S17** *A*). This rearrangement leads to a number of alterations in the relationships between H3K9me3/H3K27me3/5mC and other chromatin features in NCs: (i) H3K27me3 and H3K9me3 co-localized stronger with active marks including H3K4me1, H3K4me3, H3K27ac and RNAP II as well as H3K36me3 (**Figs. 3** *A* and **4**). (ii) 5mC co-localized somewhat stronger with H3K36me3 (**Figs. 3** *A* and **S17** *A*). (iii) Whereas 5mC and H3K27me3 were already depleted from the surface of the chromosome territory in ESCs (**Figs. 3** *C* and **S17** *B*), H3K9me3 moved into the interior of the territory in NCs (**Fig. 3** *C*). The positive correlations between H3K4me1-H3K27me3 and H3K4me1-H3K9me3 remained stronger in NCs than in ESCs also on larger genomic scales up to ten nucleosomes (**Figs. 3** *D*, **4** *C*, **S16**), indicating that they are caused by NC-specific broad domains. In summary, these findings suggest that the main chromatin transition during differentiation from ESCs into NCs is the rearrangement of H3K9me3/H3K27me3 domains, which in NCs extend beyond repressive heterochromatin and overlap at least to some extent with chromatin regions that carry activating histone marks.

**Figure 6.**
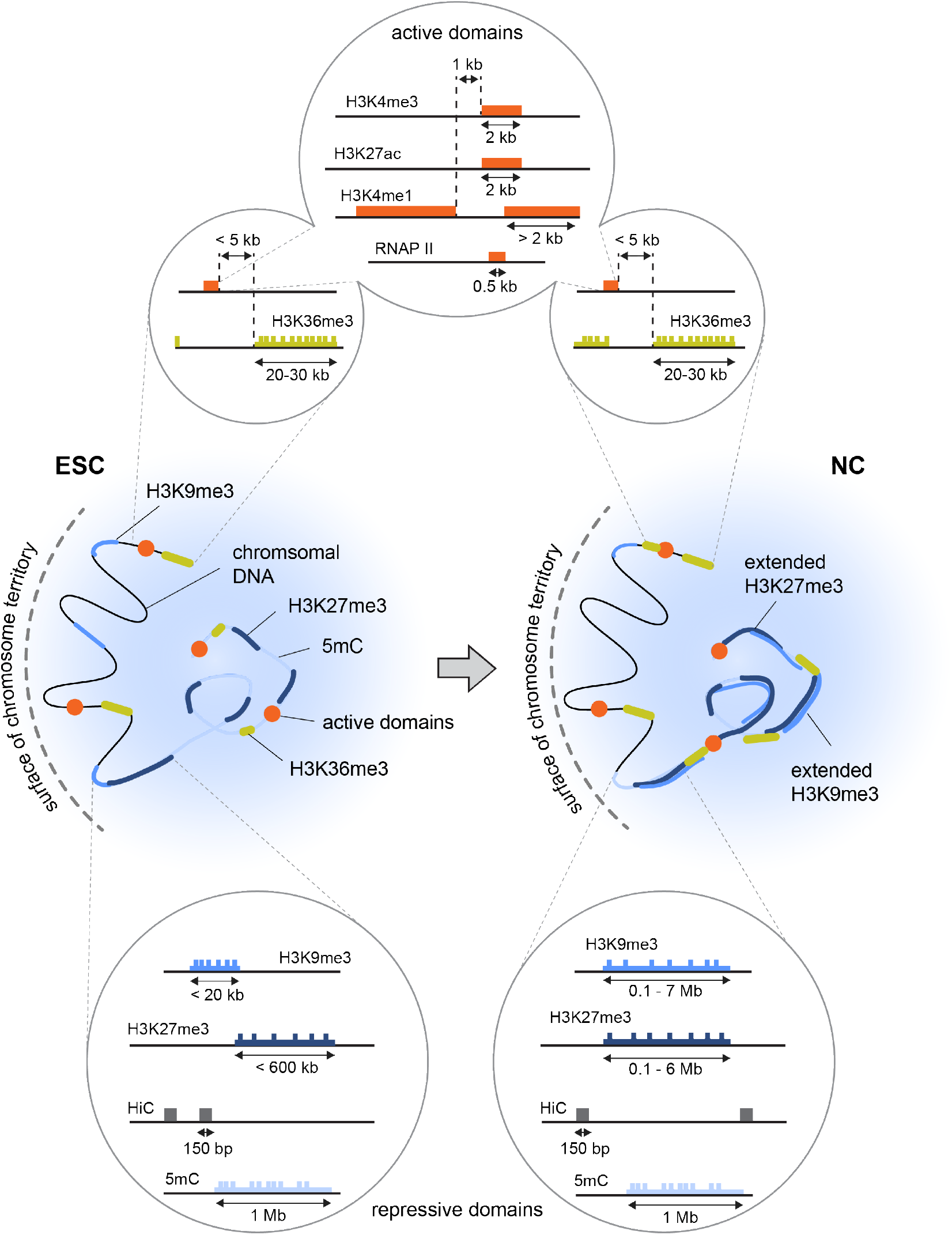
Alterations of chromatin features during differentiation of ESCs into NCs. Model for the re-organization of chromatin domains during differentiation from ESCs to NCs based on the MCORE analysis of the data sets used in this study. Active domains mostly retained their organization, with H3K4me1 being partly separated from the smaller H3K4me3/H3K27ac domains in both cell types. The overlap between those marks and H3K36me3 increased in NCs, which might be due to elevated transcription of enhancers or activation of genes enriched for H3K4me1/3 or H3K27ac. Domains enriched for H3K9me3 and H3K27me3 became extended at sites of 5mC and were preferentially buried inside chromosome territories. The newly established H3K9me3/H3K27me3 domains in NCs appeared discontinuous, i.e. contained many modified nucleosomes without an equally modified neighbor. Further, they exhibited increased overlap with activating marks such as H3K4me1 and H3K4me3, which suggests that they do not exclusively contain heterochromatin but rather enclose both active and repressive chromatin domains.

## Discussion

The quantitative understanding of how cells organize their genome into cell type-specific chromatin states is important for the description of all processes that require access to the genetic information. While the effects of soluble enzymes can be represented by simple rate equations, the polymeric nature of chromatin introduces a spatial relationship among nucleosome states. As a result, nucleosomes are influenced by the adjacent chromatin segments and patterns can form along the genomic coordinate. These patterns are present on different length scales and represent an extra layer of complexity, which is an essential part of the regulatory networks that control genome functions. For example, repressive histone modifications form broad domains that are relatively independent from the underlying DNA sequence and can be transmitted through at least several cell divisions (22, 53–55). Furthermore, chromosomes fold into topological domains that determine the contact frequencies between genomic loci and the proteins they are decorated with (56), thereby creating three-dimensional structural patterns that might be relevant for long-range gene regulation. Elucidating the mechanistic basis of these phenomena and the functional relationships among them requires techniques that can identify, quantitate and compare different patterns along the genome.

### Global analysis of deep sequencing data by correlation functions

The analysis of deep sequencing data on the level of individual genomic positions is complicated by noise, bias and undersampling (15-17). It is often not straightforward to choose a threshold value for classifying enriched regions because low values lead to false-positive peaks and high values lead to false-negative results. Consequently, identifying differences in the chromatin domain landscape between samples is currently fraught with difficulties, which is evident from a comparison of 14 different software tools for differential ChIP-seq analysis that yield different results (57). These problems are especially detrimental for the analysis of broad regions with low enrichment levels that are common to heterochromatin.

The MCORE method introduced here uses correlation functions to find and quantify chromatin patterns. It computes Pearson correlation coefficients as underlying metrics, which is a convenient measure that has extensively been used for data comparison and statistical inference in many fields including deep sequencing analysis (18, 30, 31, 58).

When calculating correlation functions, MCORE implicitly combines multiple genomic regions to gain a correlation coefficient for each shift distance, yielding statistical robustness from a large number of reads. In this manner MCORE can quickly retrieve information on the spatial distribution of chromatin features on all length scales, while avoiding assumptions or model-dependent parameter settings like significance thresholds. In contrast to aggregate plots (59-61) MCORE does not rely on any *a priori* knowledge about annotated genomic elements. Compared to peak calling (15), MCORE has a relatively low sensitivity to undersampling. This might be beneficial for the analysis of data sets that have low complexity, e.g. due to limitations in input material as it is the case for low input sequencing samples, or insufficient sequencing depth, which seems to be the norm for broadly distributed histone modifications (15). Domain abundances obtained from data sets with different coverage values exhibited somewhat larger changes than domain sizes. Therefore, sufficient coverage should be ensured in order to interpret these parameters, e.g. by applying MCORE to diluted data as shown in **Figs. S5** and **S6**.

A crucial step in the MCORE workflow is correction for bias and background. Without this step artificially overrepresented regions and non-specific signal can induce similarities between data sets that are unrelated to the chromatin feature of interest. These phenomena are well known from other deep sequencing analysis methods. Because different artifacts affect the signal on different scales, their contribution and successful correction can better be assessed by multi-scale methods than by techniques that operate on a single scale. In particular, non-specific background leads to a characteristic correlation spectrum whose removal can and should be validated using the proper controls. Based on a single correlation coefficient between data sets this task is more difficult to accomplish. Occupancy profiles that have been normalized according to the workflow presented here might serve as a useful resource for other downstream analysis methods.

### Genome-wide topology of chromatin domains

MCORE extends previous techniques that assess co-localizations of chromatin features based on correlation coefficients. By evaluating entire correlation functions instead of single correlation coefficients the spatial extension of chromatin patterns on multiple genomic scales is retrieved. With this analysis we found predominantly small domain sizes of less than 2 kb for promoter/enhancer marks H3K4me1, H3K4me3, H3K27ac and RNAP II, intermediate domain sizes of 20–30 kb for H3K36me3 that marks the whole gene body including flanking regions, and domain sizes up to several megabases for H3K9me3/H3K27me3. This is consistent with the size of promoters, enhancers and active genes, and with the estimates for repressive domains that were made based on visual inspection of selected genomic regions (62).

The scale-dependent relationships determined by MCORE for different histone modifications suggest that there are three types of domain topologies: (i) Short domains formed by activating marks are relatively homogenously modified, which is reflected by a large probability for finding the same or another activating modification at the next nucleosome. Accordingly, correlation functions for activating marks such as H3K4me3 displayed only a moderate initial decay (**Fig. 2**). (ii) H3K36me3 formed domains of intermediate size that were 1–2 orders of magnitude broader than H3K4me3 domains. The stronger initial decay (**Fig. 2**) suggests the presence of single nucleosomes without an equally modified neighbor, which is consistent with the presence of more gaps in H3K36me3 domains as compared to H3K4me3 domains. (iii) Especially in NCs, replicate correlation functions for H3K9me3 or H3K27me3 displayed long-range correlations that extended to shift distances of several megabases. Similar scale-dependence was also seen for correlation functions between H3K9me3 and H3K27me3 (**Fig. 3** *C*), suggesting that these domains are intermingled. The respective correlation functions displayed a relatively fast decay at a shift distance of one nucleosome (**Figs. 2** and **3**), indicating that many modified nucleosomes within these broad domains localize next to a non-modified or differently modified one. Such a domain structure fits well to the experimental observation of broad domains and low enrichment levels in the cell ensemble. In particular, the experimentally determined methylation levels that are below
50% even for H3K9me3 in pericentric heterochromatin (see (22) and references therein) are incompatible with large genomic regions containing exclusively fully H3K9me3-modified nucleosomes. Broad H3K9me3/H3K27me3 domains with gaps are consistent with a model in which methylation marks are stochastically propagated from well-positioned nucleation sites via dynamic chromatin looping (22, 63).

### Comparison of chromatin domains in ESCs versus NCs

The comparative analysis of 11 different chromatin features in ESCs and NCs conducted here shows that MCORE can efficiently identify and compare chromatin domain patterns. By integrating genome-wide data sets with very different readouts MCORE is well suited to generate hypotheses that can be further validated in downstream applications.

The positive correlations we found among activating histone modifications (H3K4me1, H3K4me3, H3K27ac, H3K36me3), among repressive histone modifications (H3K9me3, H3K27me3, 5mC) and between H3K36me3 and repressive marks are in qualitative agreement with previous studies conducted with ESCs and other cell types (62, 64, 65). Genome-wide co-localization of marks that were originally thought to affect transcription antagonistically might reflect the additional functions of these marks that are unrelated to the regulation of gene expression. For example, H3K9me3 is not restricted to heterochromatin but is also found at active genes (47, 66). Furthermore, H3K9me3, H3K27me3 and H3K36me3 have been linked to alternative splicing (51, 67) and large portions of H3K9me3 and H3K27me3 localize to intergenic regions where they might serve completely different functions (64). Because sequencing data reflect the average of the cell population that was analyzed, positive correlations might also arise from gene loci carrying different marks during different cell cycle stages, alleles within the same cell carrying different marks, or loci carrying different marks in different cells. The finding that correlations were generally smaller in ESCs than in NCs fits to the model of plastic and ‘hyperactive’ chromatin in stem cells, which acquires distinct patterns only upon differentiation (68). The fact that most 5mC regions persisted in ESCs and NCs, were moderately depleted for inter-chromosomal contacts in both cell types, and gained H3K9me3 in NCs suggests a model in which heterochromatic regions newly established in NCs are preferentially buried within chromosome territories (**Fig. 6**). H3K27me3 domains behaved similarly in both cell types, which fits very well to the previously reported localization of inactive domains such as the Hox cluster inside chromosome territories in differentiated cells (13, 69–71). The observation that only a subset of H3K9me3 domains is broad and enriched for SUV39H1 suggests that heterochromatin extension is not merely caused by recruitment of frans-acting enzymes to preexisting H3K9me3 but rather by site-specific recruitment of methyltransferases to domains that are to be extended during differentiation. Although further experiments are required to fully understand the underlying molecular details of heterochromatin reorganization during differentiation, these insights provide a starting point to uncover the pathways that are responsible for establishing differently sized heterochromatin domains with distinct molecular composition.

## Conclusions

The MCORE method introduced here enables the quantitative retrieval and comparison of patterns and spatial relationships for different chromatin features from noisy data sets. These features make MCORE complementary to model-dependent approaches that assess the local read density at individual loci to find enriched regions. MCORE is relatively fast and yields a coarse-grained comparison of data sets without the requirement of user-defined input parameters, providing an unbiased starting point for in-depth analyses conducted downstream. We anticipate that MCORE will aid in the design and validation of mechanistic models for chromatin patterning and long-range gene regulation.

## Author contributions

FE and KR designed research. JM and FE performed the theoretical work. All authors analyzed and interpreted the data. JM and JPM performed the experiments. The manuscript was written by FE, JM and KR.

## Acknowledgments

We thank Caroline Bauer for valuable assistance, the DKFZ Genomics and Proteomics Core Facility for technical support and expertise, and Anne Rademacher, Katharina Müller-Ott and Daniel Duzdevich for comments on the manuscript. This work was supported by grant CA146 of the Cancer Research Cooperation Program between the DKFZ and the Israel Ministry of Science and Technology (MOST) and the projects ImmunoQuant (0316170B) and PRECiSe (031L0076A) of the German Federal Ministry of Education and Research (BMBF) as well as a DKFZ intramural grant to FE.

## Supporting Citations

References (72-100) appear in the Supporting Material.

## Supporting Material

### Supporting Figures

1. Strategies to retrieve information about complex patterns
2. MCORE workflow and background correction
3. Statistics and Spearman correlation functions for representative ChIP-seq data
4. Peak calling for representative ChIP-seq data
5. Robustness of correlation functions towards undersampling
6. Dependence of fit results on coverage
7. MCORE for simulated data sets
8. Errors and statistical comparison of correlation functions
9. MCORE for different H3K27ac data sets
10. Quality control of ChIP-seq data
11. Fitted correlation functions for H3K27me3
12. Peak calling summary for H3K9me3
13. MCORE-directed annotation of chromatin features
14. MCORE for transcription factor binding
15. Spatial extension and co-localization of different features in ESCs versus NCs
16. Heterochromatin reorganization during differentiation
17. DNA methylation and inter-chromosomal contacts

### Supporting Tables

1. Comparison of MCORE with other software tools
2. Overview of chromatin features assessed in this study
3. Fit parameters for selected correlation functions in ESCs
4. Fit parameters for selected correlation functions in NCs
5. Summary of data sets used in this study

**Figure S1.**
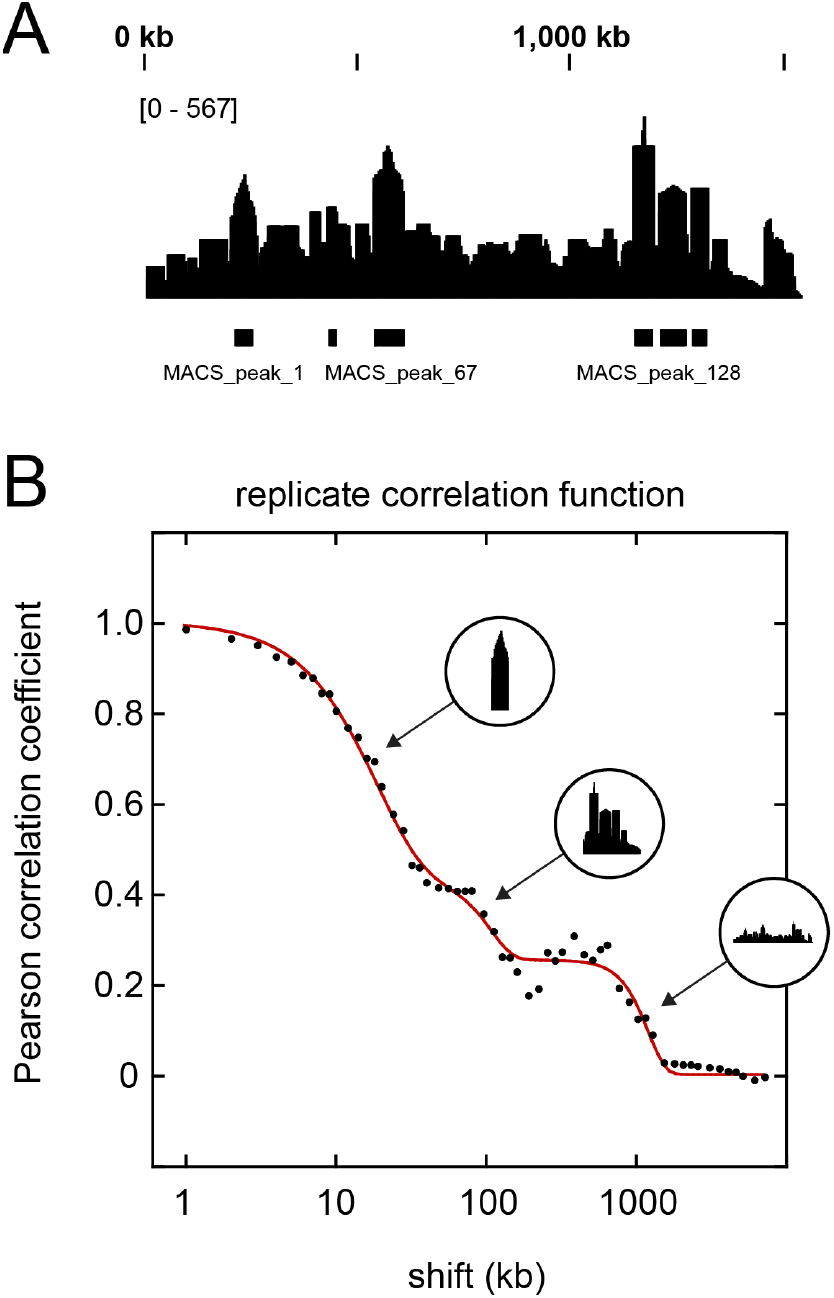
Strategies to retrieve information about complex patterns. **(A)**Peak calling result for a complex domain structure involving different enrichment levels (MACS, standard settings mfold =10,30, pvalue = 1e-5). The pattern is reduced to regions that are compatible with the threshold and significance settings while others are ignored. **(B)**Correlation function (black dots) and multi-exponential fit according to Eq. 6 (red line) for the pattern in panel A. The correlation function yields the different length scales that are present in the pattern, including the width of highly enriched regions, the characteristic size of clusters formed by adjacent peaks, and the size of the entire enriched region.

**Figure S2.**
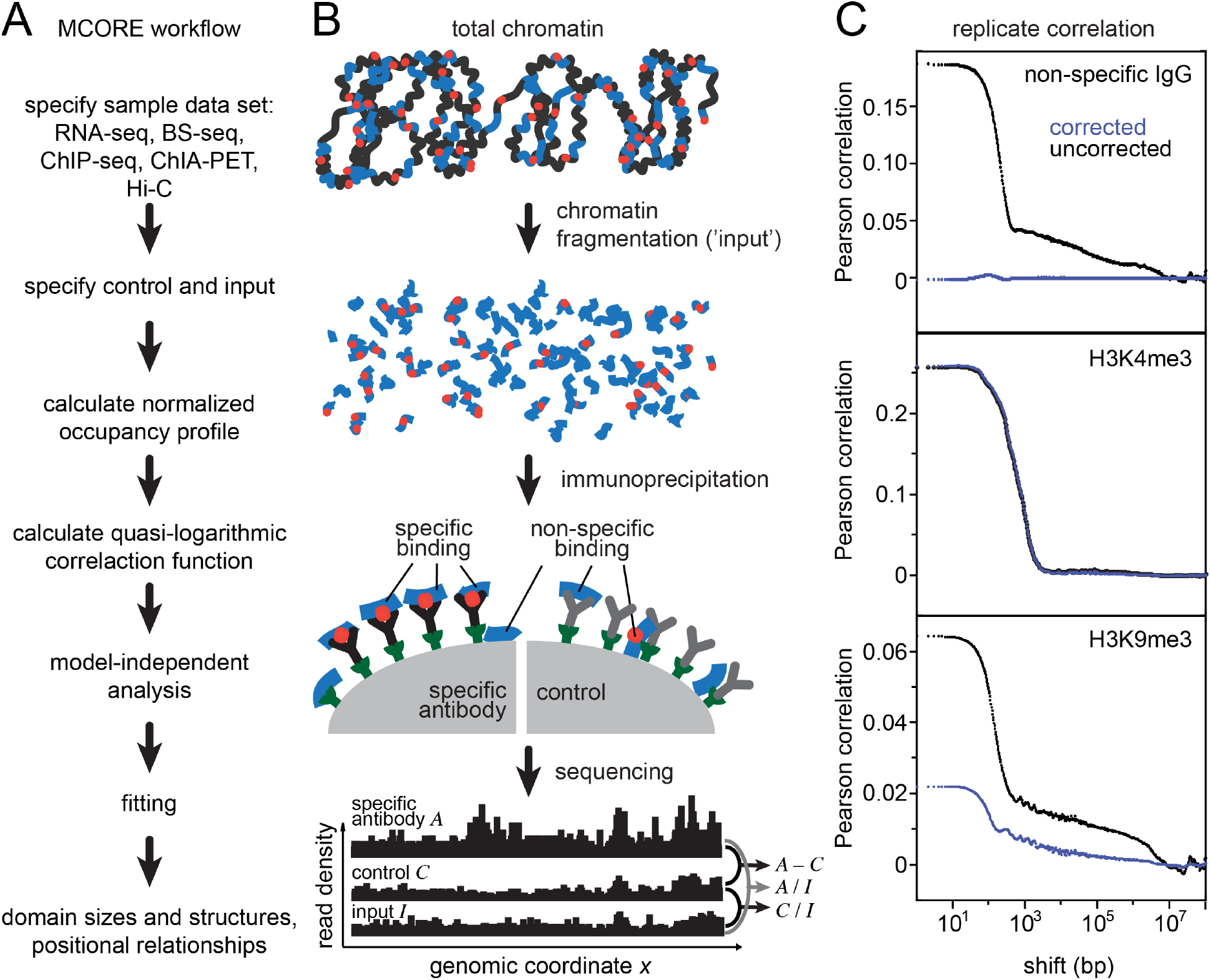
MCORE workflow and background correction. **(A)** Schematic representation of the MCORE workflow. **(B)**Fragmentation of total chromatin (black) containing a chromatin feature of interest (red) occurs with some bias and is frequently incomplete. As a result, only a fraction of chromatin (blue) is present in the input sample due to size selection during library preparation. Subsequent immunoprecipitation occurs in the presence of non-specific binding. The latter contribution can be assessed in a separate control reaction, e.g. by using an antibody that does not bind specifically to the antigen. Sequencing reads obtained from samples with the specific antibody *A*, the control *C* and the input *I* are used to calculate normalized occupancy profiles for the analysis of a given chromatin feature according to Eqs. 1–3. In brief, the read densities from the specific IP and from the control are divided by the input density (*A / I* and *C/I*, see Eq. 1) to account for multiplicative biases such as mappability or preferences in immunoprecipitation, ligation, amplification and sequencing. Next, the weighted control signal is subtracted from the specific antibody signal to remove additive bias caused by non-specific binding (Eqs. 2-3). Resulting profiles are used for calculating correlation functions (Eq. 4). **(C)**Correlation functions for the uncorrected (black) and corrected (blue) occupancies for control IP (IgG, top), H3K4me3 (center) and H3K9me3 (bottom) ChIP-seq replicates in neural progenitor cells. Subtraction of the weighted control IP signal removes the background correlation and thus eliminates correlation between control IP signals (top). Normalization has little effect for H3K4me3, which displays distinct peaks with considerable enrichment (**Fig. S4**). In contrast, it causes a significant correction for H3K9me3, which forms broad domains with moderate enrichment levels.

**Figure S3.**
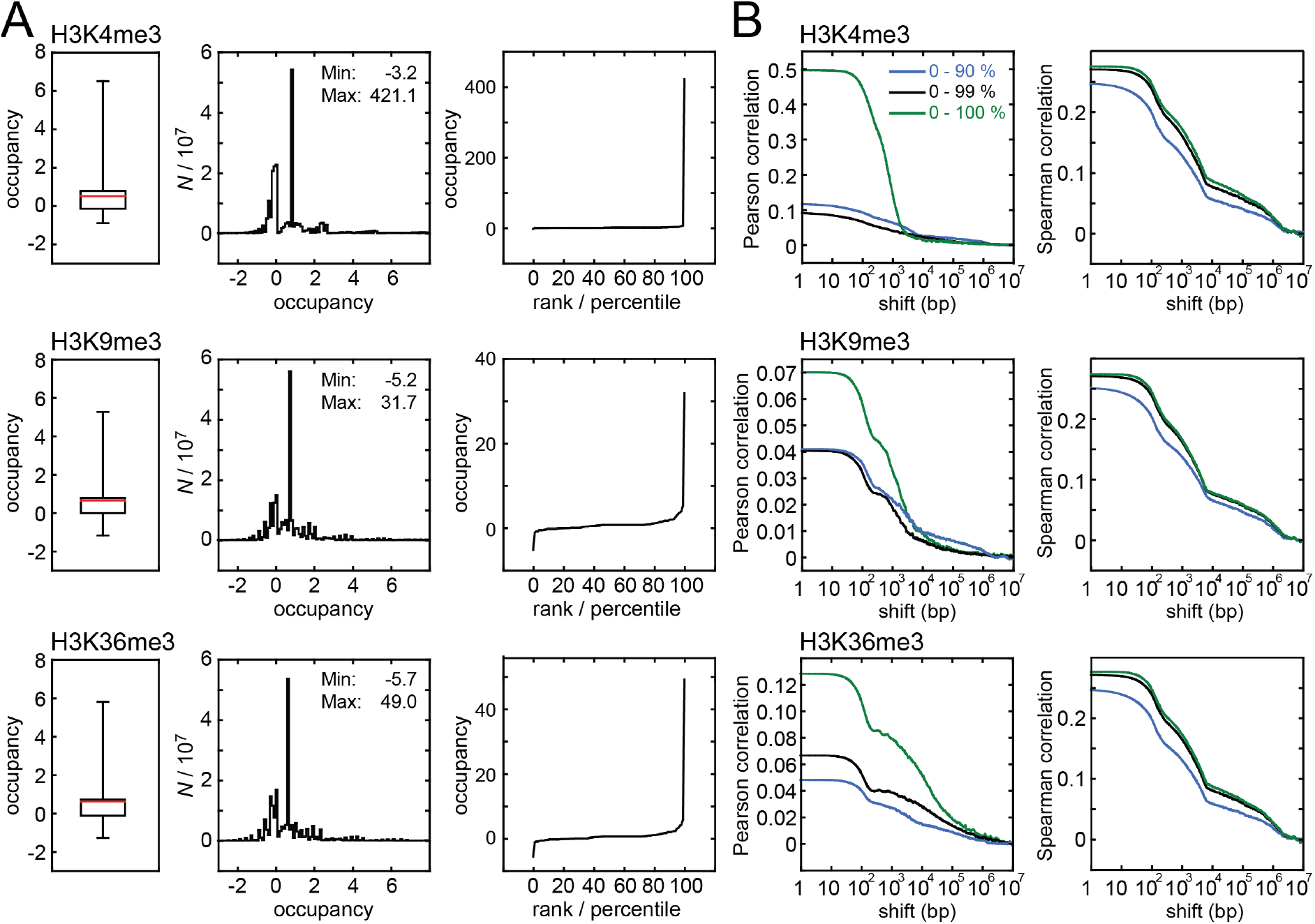
Statistics and Spearman correlation functions for representative ChIP-seq data. **(A)** Box plots (left), histograms (center) and percentiles (right) for normalized occupancy profiles from H3K4me3, H3K9me3 and H3K36me3 ChIP-seq experiments in ESCs. For box plots, the median is colored in red and the ends of the whiskers represent the 1^st^ and 99^th^ percentile. Minimum and maximum occupancy values are listed in the histograms. The background comprises a large part of the data and its distribution is similar for all profiles (see box plots and histograms). **(B)**Pearson (left, green) and Spearman (right, green) correlation functions for the occupancy profiles analyzed in panel *A*. To assess the contribution of enriched regions to the different correlation functions we replaced occupancy values above the 90^th^ (blue) or 99^th^ (black) percentile with the average occupancy within the rest of the genome. Spearman correlation functions exhibited only slight changes upon removal of highly enriched regions and primarily reflected the structure of the background signal that was independent of the interrogated histone mark (compare top, center and bottom in the right column). In contrast, Pearson correlation functions reflected the properties of enriched regions, which carry the biological information, and changed their shape when these regions were omitted from the analysis. After removal of enriched regions (left column, blue), Pearson correlation functions were dominated by the background signal and resembled Spearman correlation functions (right column). The stronger background signal in Spearman correlation functions is due to the correction procedure that minimizes the background according to the Pearson metric (Eq. 3).

**Figure S4.**
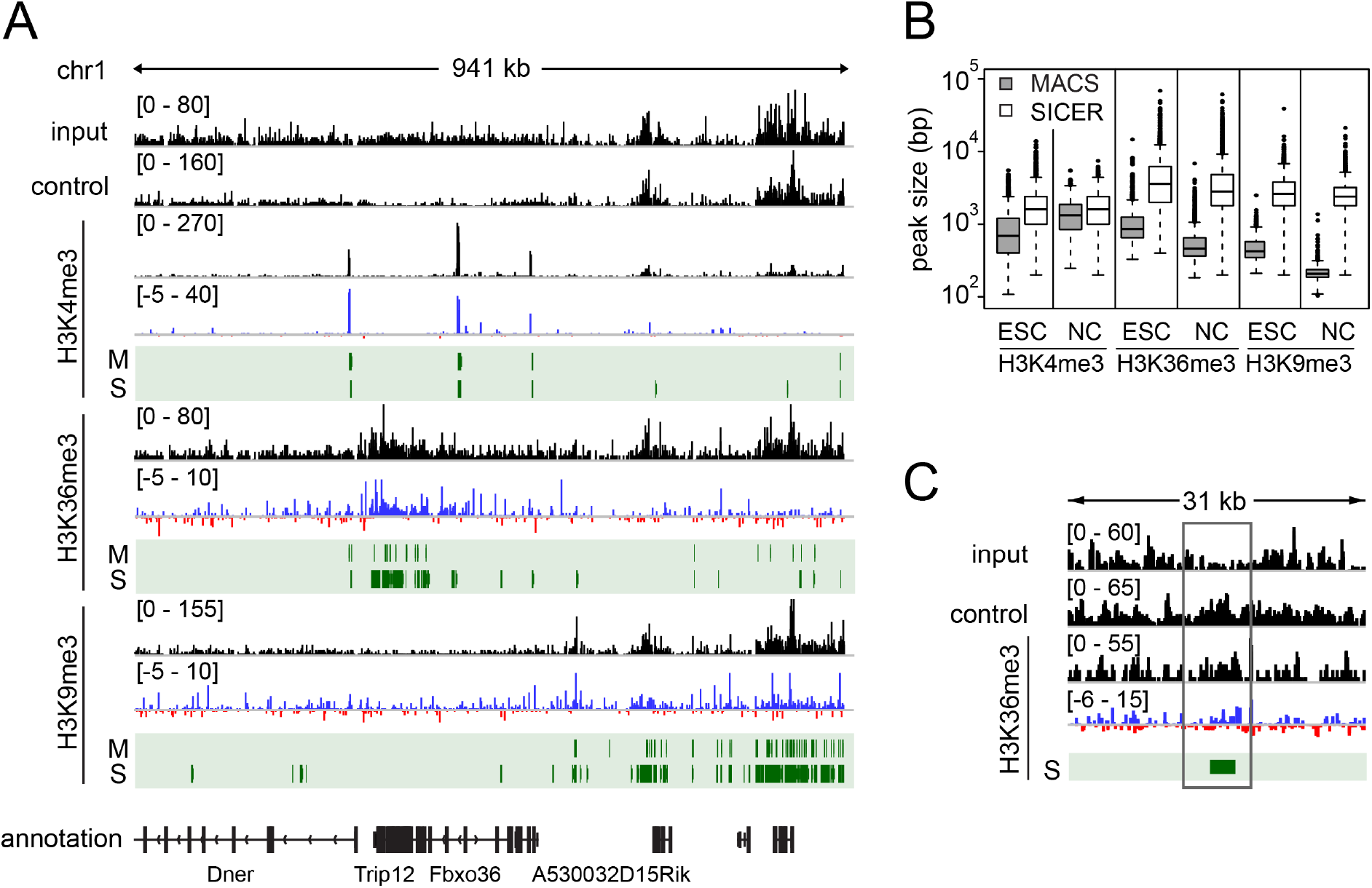
Peak calling for representative ChIP-seq data. **(A)** Read distribution (black) for sample, control (IP with a non-specific antibody) and input, normalized occupancy (red/blue), and peaks (green) called by MACS (M) and SICER (S) for H3K4me3, H3K9me3 and H3K36me3 ChIP-seq in NCs. Distinct H3K4me3 domains were reliably identified by both peak callers, results for H3K9me3 and H3K36me3 depended on the specific algorithm used (e.g. MACS and SICER). **(B)** Peak size distributions for clusters called by MACS and SICER for the ChIP-seq experiments in ESCs and NCs. Resulting cluster sizes differed between both methods. **(C)** Example of the read distribution (black) and normalized occupancy (red/blue) for H3K36me3 ChIP-seq in NCs, including input and control. The highlighted region contains an apparent enrichment in H3K36me3 that is identified as a peak. However, similar enrichment is present in the control IP, suggesting that the signal corresponds to non-specific background.

**Figure S5.**
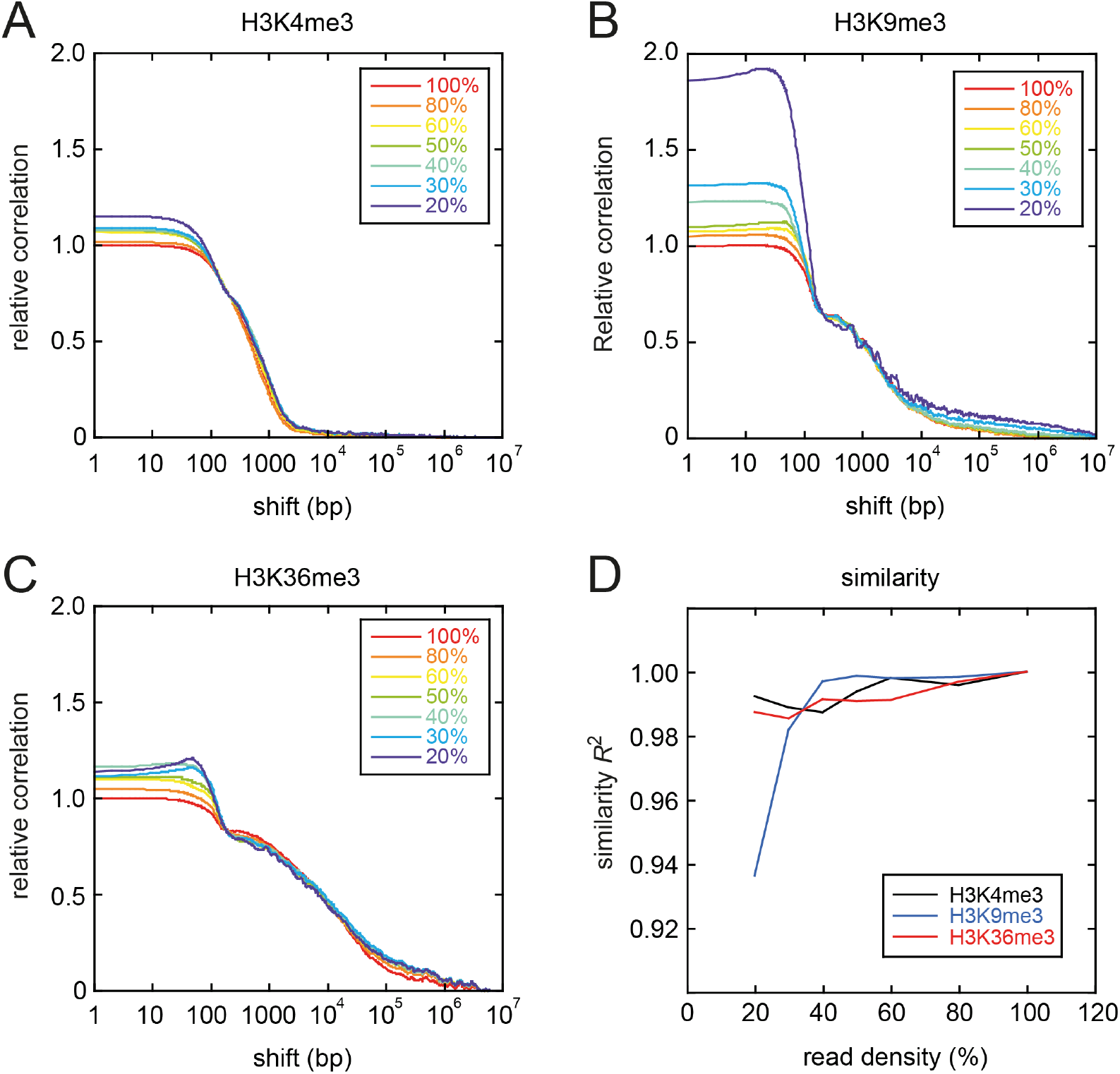
Robustness of correlation functions towards undersampling. **(A)**Replicate correlation functions for ChIP-seq data sets of H3K4me3 in ESCs containing different numbers of reads. The red curve corresponds to the entire set of reads reported in this study (100%, corresponding to 30 million reads). The other functions reflect data sets that were diluted *in silico* by randomly selecting only a fraction of reads from the entire set. Correlation functions were normalized to the 100% curve at a shift distance of one nucleosome(according to the fit parameters c2 in **Table S3**) because correlation coefficients for smaller shift distances do not contain information about domain structures (see **Fig. S6**for domain sizes obtained by fitting). **(B)** Same as in panel *A* but forH3K9me3. **(C)**Same as in panel *A* but for H3K36me3. **(D)** Quantification of the similarity of correlation functions for diluted data sets with respect to the curve for the undiluted data set based on the coefficient of determination (*R*^2^). Correlation functions for diluted data sets are similar to each other and to the result forthe undiluted data set, with *R*^2^> 0.9. Above 40% read density, which corresponds to 12 million reads, a plateau is reached for all modifications assessed here.

**Figure S6.**
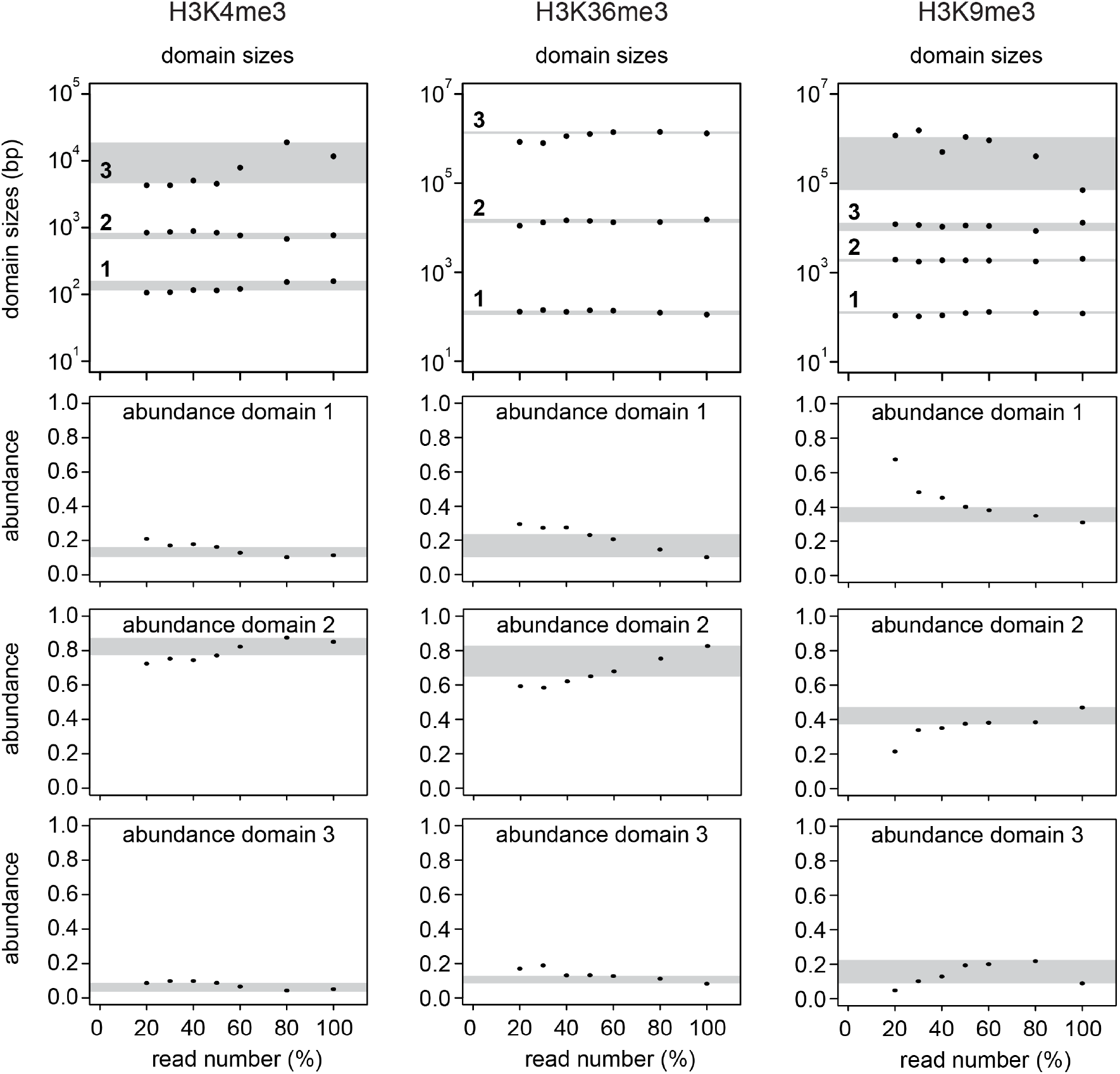
Dependence of fit results on coverage. The correlation curves plotted in **Fig. S5** were fitted with Eq. 6. Fit results for the domain sizes and the respective amplitudes are plotted versus coverage (domain numbers are indicated in the top panel). Gray regions show the variation of the fit results for dilution down to 50% of the reads. The most abundant domains, which represent the characteristic domain sizes for a given modification, were accurately quantified from diluted functions (top panels). Only lowly abundant large domains like the largest domain for H3K4me3 or H3K9me3 with abundance below 10% (see **Table S3**for values) changed their apparent size when coverage was reduced. Due to their low abundance we did not interpret these domains.

**Figure S7.**
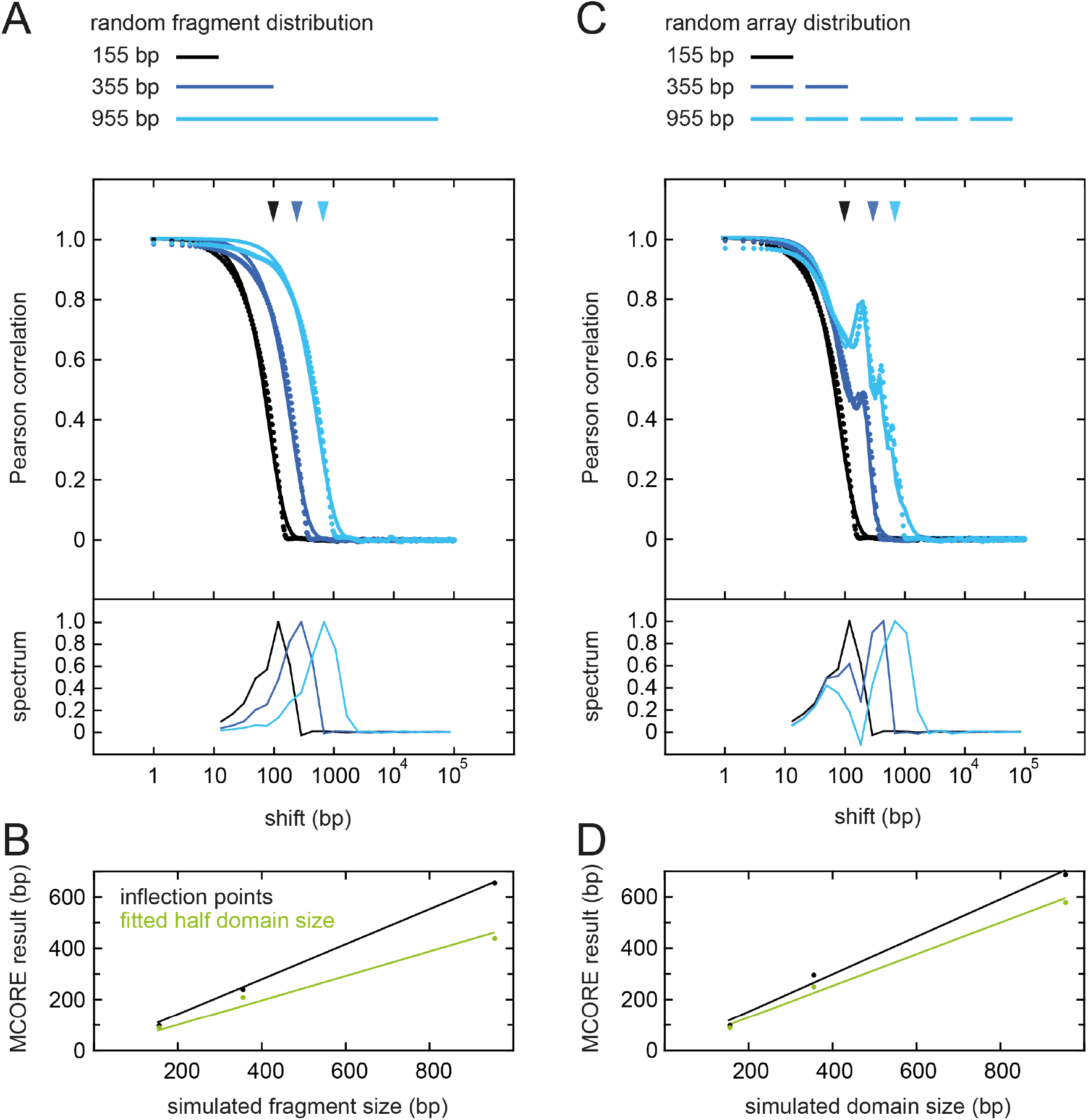
MCORE for simulated data sets. **(A)** Correlation functions (dotted lines) for randomly distributed fragments of different size exhibit a single decay length that can be retrieved by assessing inflection points (arrowheads), by fitting the model function in Eq. 7 (solid lines) or by evaluating the decay spectrum obtained from the Gardner transformation shown below the curves. **(B)** Fit parameters obtained for the curves shown in panel *A* yield half domain sizes (green), whereas the positions of inflection points correspond to 0.7-times the domain sizes (black). **(C)** Correlation functions (dotted lines) for nucleosomal arrays (instead of continuous fragments as in panel *A*) display global decay lengths that correspond to array sizes. The decay lengths coincide with the largest inflection points depicted by the arrowheads. In addition, correlation functions exhibit an oscillatory contribution due to the nucleosomal pattern within the arrays. The nucleosome repeat length of 200 bp used for the simulation was retrieved by fitting with Eq. 7 (solid lines). **(D)** The array size in panel *C* is either obtained from the analysis of inflection points (black), the peaks of the decay spectrum or the fitted half domain sizes (green), with the same scaling found for continuous domains in panel **B**.

**Figure S8.**
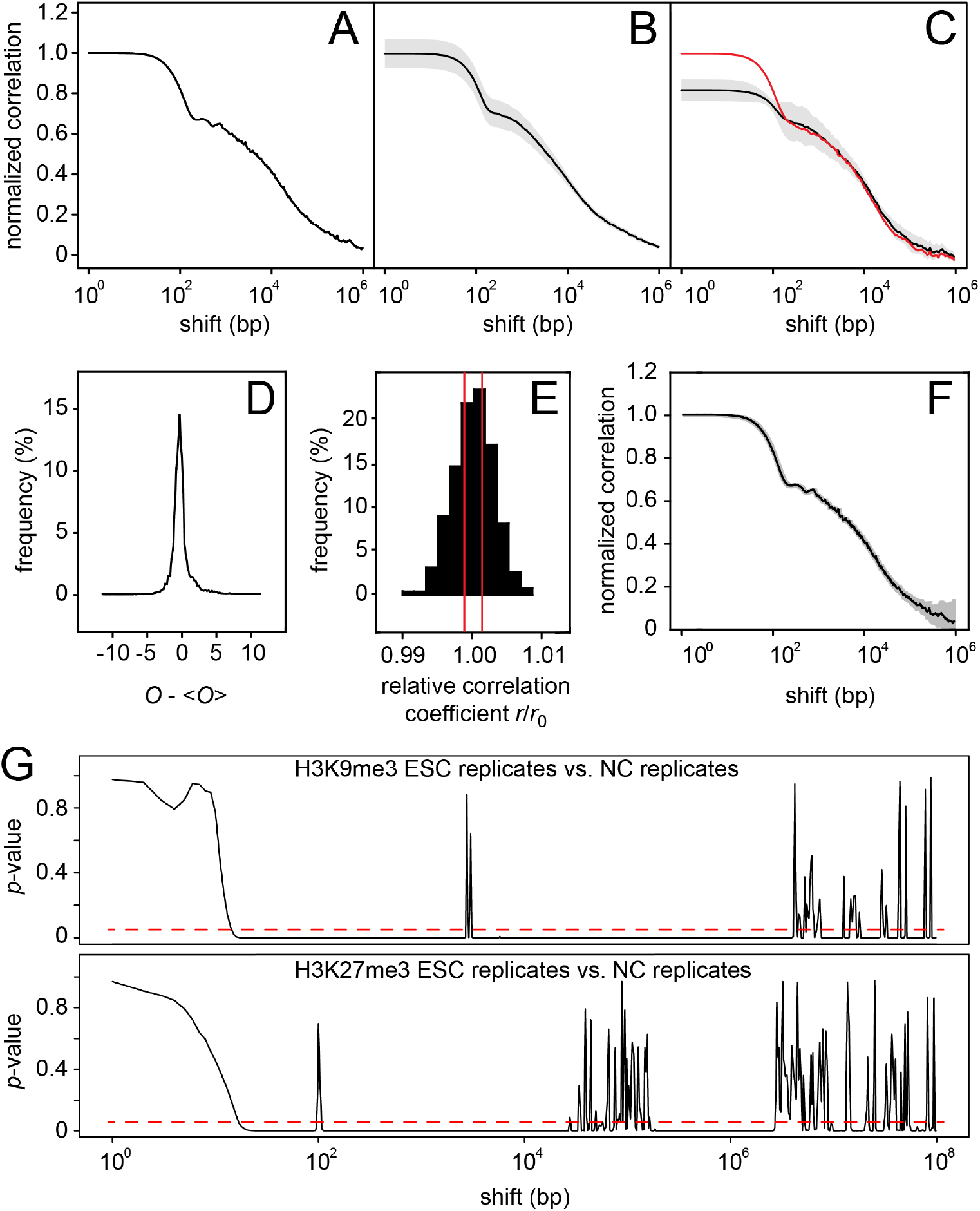
Errors and statistical comparison of correlation functions. Panels *A-F* refer to H3K36me3 in ESCs. **(A)** Replicate correlation function (black) and its confidence interval (gray) obtained using the Fisher transformation (Materials and Methods). Due to the large sample size the confidence interval is smaller than 10^-3^ and within the line thickness. **(B)** Average (black) and confidence interval (gray) of correlation functions calculated for all autosomes (1-17) based on the H3K36me3 data sets generated in this study. **(C)** Average (black) and confidence interval (gray) of three replicate correlation functions calculated from three independent biological replicates (rep1 x rep2, rep1 x rep3, rep2 x rep3), yielding information on experimental reproducibility. The correlation function for ENCODE data for H3K36me3 in ESCs (red) is similar to the correlation function computed from the data sets generated in this study. The amplitude of the first domain that covers the length scale below 200 bp shift distance is different. This might be due to incomplete correction of background signal in the ENCODE data set that lacks a control IP reference, which should, however, not strongly affect the quantitation of domain sizes beyond the scale of a nucleosome. **(D)** Distribution of normalized occupancy values (*O_i_ - >O<*) that were used for calculating the correlation function in panel *A*. The distribution is relatively symmetric and unimodal. **(E)** Distribution of correlation coefficients obtained by bootstrapping for the correlation coefficient at zero shift distance. Each correlation coefficient was calculated after resampling the occupancy profiles with replacement as described in the Materials and Methods section. Correlation coefficients are given relative to the mean value. The 95% confidence interval obtained by this approach is roughly 3-times larger than the estimate based on Fisher transformation (shown in red). **(F)** Correlation function from panel *A* with non-parametric bootstrap confidence intervals for each shift distance. **(G)** Based on 95% confidence intervals, the statistical significance of differences between correlation functions can be assessed. *p*-values for the difference of two functions at each shift distance are shown, which were calculated based on a *t*-test for each pair of correlation coefficients. The red dashed lines indicate a *p*-value of 0.05. Top: Comparison between H3K9me3 in ESCs and NCs. Correlation curves are shown in **Fig. 3** *B* (top, black/blue). Bottom: Comparison between H3K27me3 in ESCs and NCs. Correlation functions are shown in **Fig. 3** *B* (center, black/blue).

**Figure S9.**
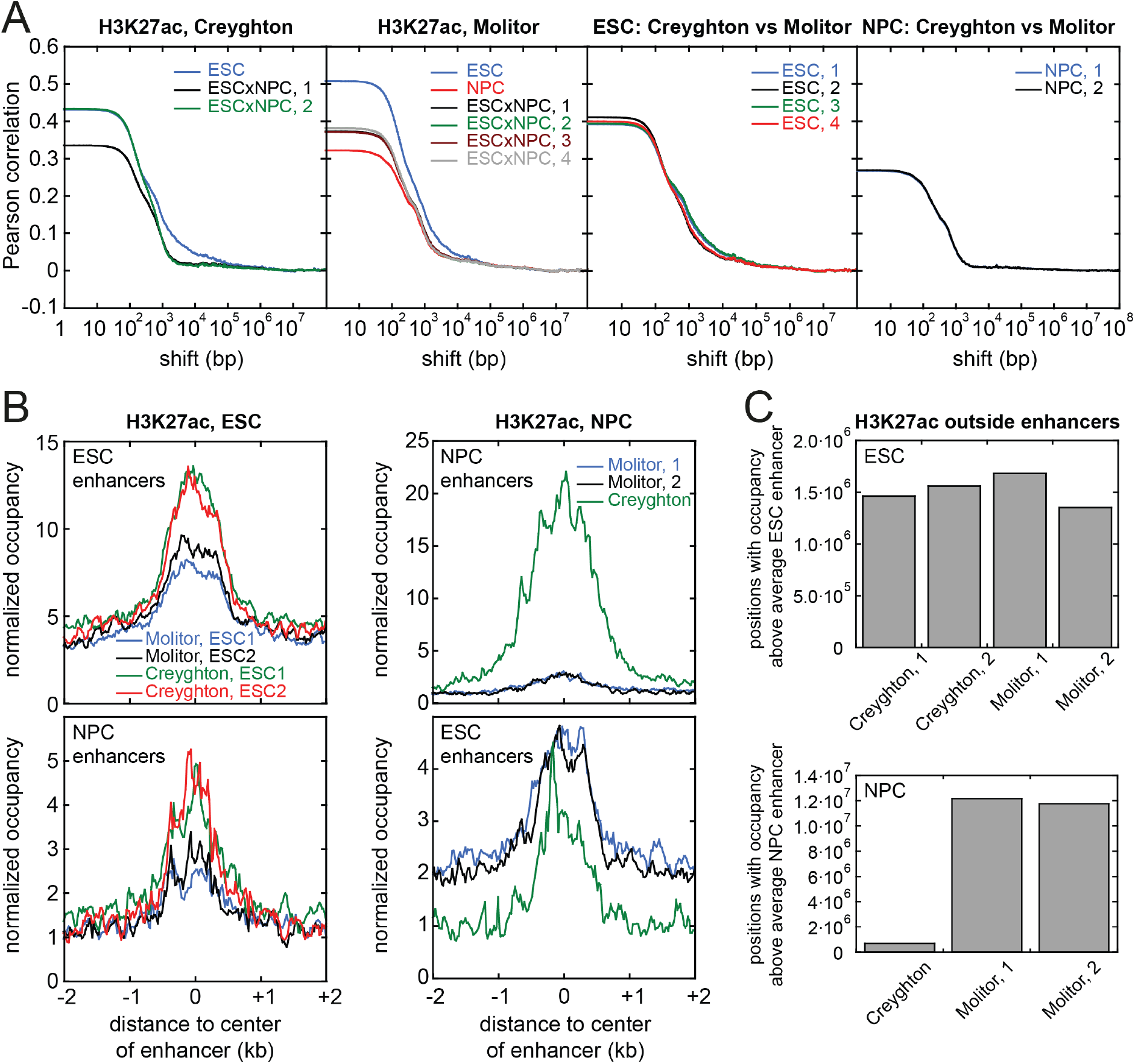
MCORE for different H3K27ac data sets. **(A)** Correlation functions for H3K27ac data sets from this manuscript (‘Molitor’) and from the study of Creyghton et al. (1) (‘Creyghton’). Both data sets yielded similar results in the MCORE analysis. **(B)** H3K27ac enrichment at the enhancers identified by Creyghton et al. was found for all data sets assessed here. **(C)** The enhancers identified by Creyghton et al. were not the only genomic regions enriched for H3K27ac.

**Figure S10.**
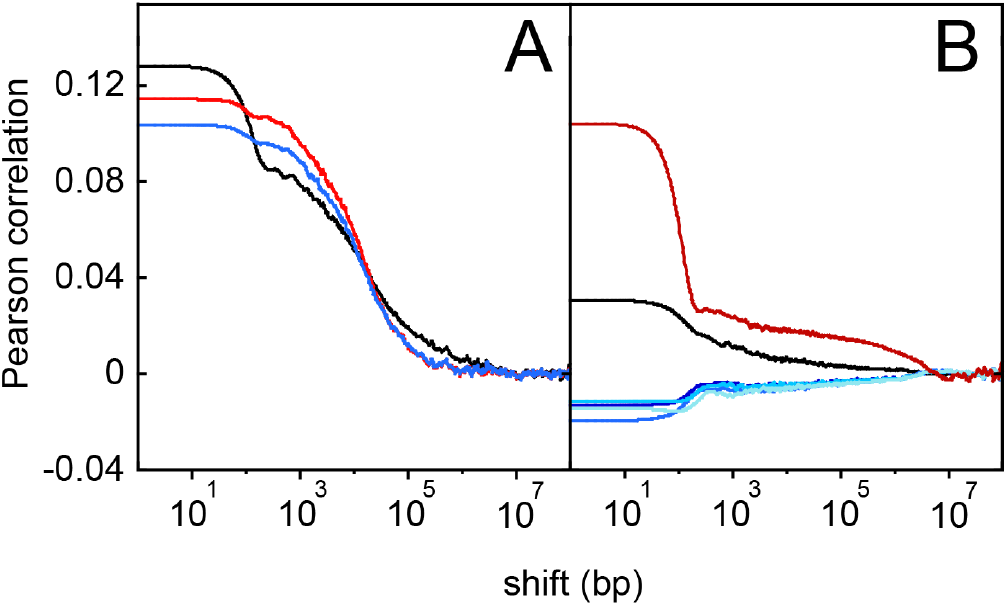
Quality control of ChIP-seq data. **(A)** Replicate correlation functions from three ChIP-seq experiments of H3K36me3 in ESCs for all pairwise combinations, replicate 1 and 2 (black), replicate 1 and 3 (red), replicate 2 and 3 (blue). The correlation functions show variations that reflect the biological reproducibility of the experiment. **(B)** Evaluation of two different antibodies used for ChIP-seq of H3K9ac in ESCs. Two ChIP-seq experiments were conducted with polyclonal antibodies from abcam (ab4441, replicate ab1 and ab2) or Active Motif (#39137, replicates am1 and am2). Replicate correlation functions of experiments with the same antibody showed significant correlation (ab1 and ab2, red line; am1 and am2 black line) with a difference in the amplitude that indicates a higher similarity and therefore a better reproducibility of ChIP-experiments conducted with the Abcam antibody. Cross-correlation functions calculated for data sets using different antibodies (blue curves for every combination of two replicates, ab1 x am1, ab1 x am2, ab2 x am1, ab2 x am2) yielded negative correlations. Thus, the two antibodies recognize different chromatin features and further validation is necessary to make conclusions on the H3K9ac distribution.

**Figure S11.**
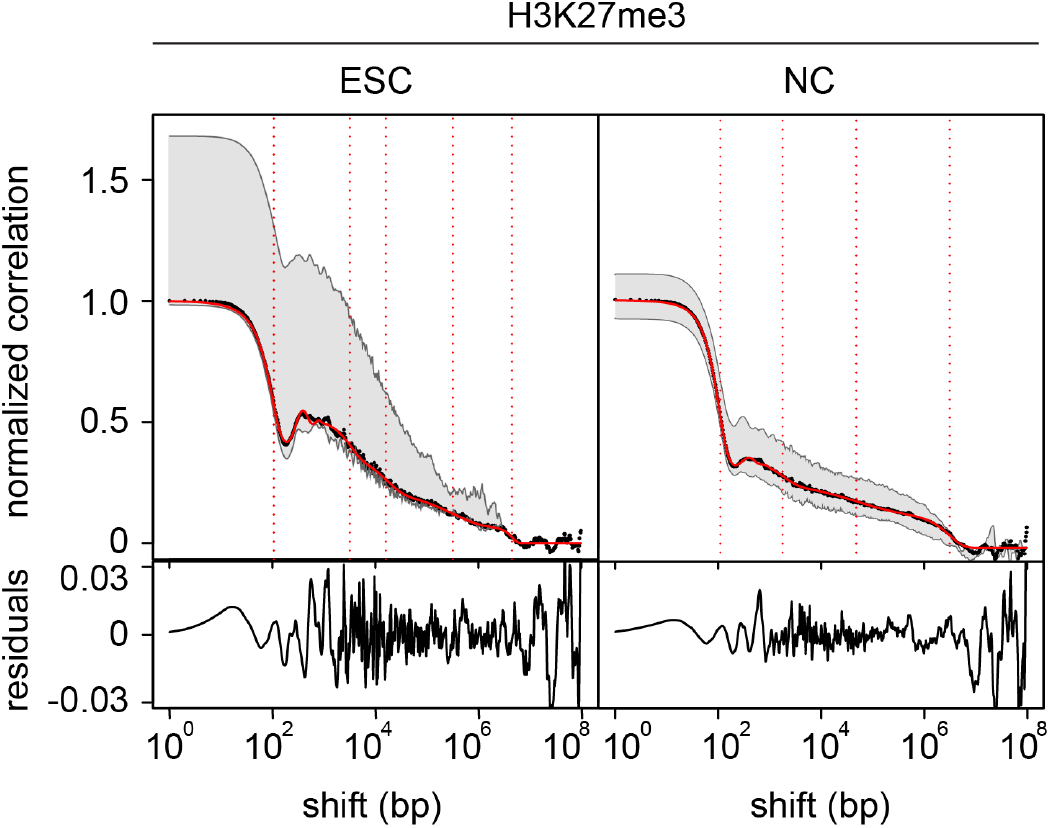
Fitted correlation functions for H3K27me3. Correlation functions calculated between replicates on chromosome 1 (black) and fit functions according to Eq. 7 (red) with half domain sizes obtained from the fit (vertical dotted lines). Gray regions indicate maximum variation among chromosomes. Fit residuals for the correlation functions are shown below the curves. Fit parameters are summarized in **Tables S3** and **S4**.

**Figure S12.**
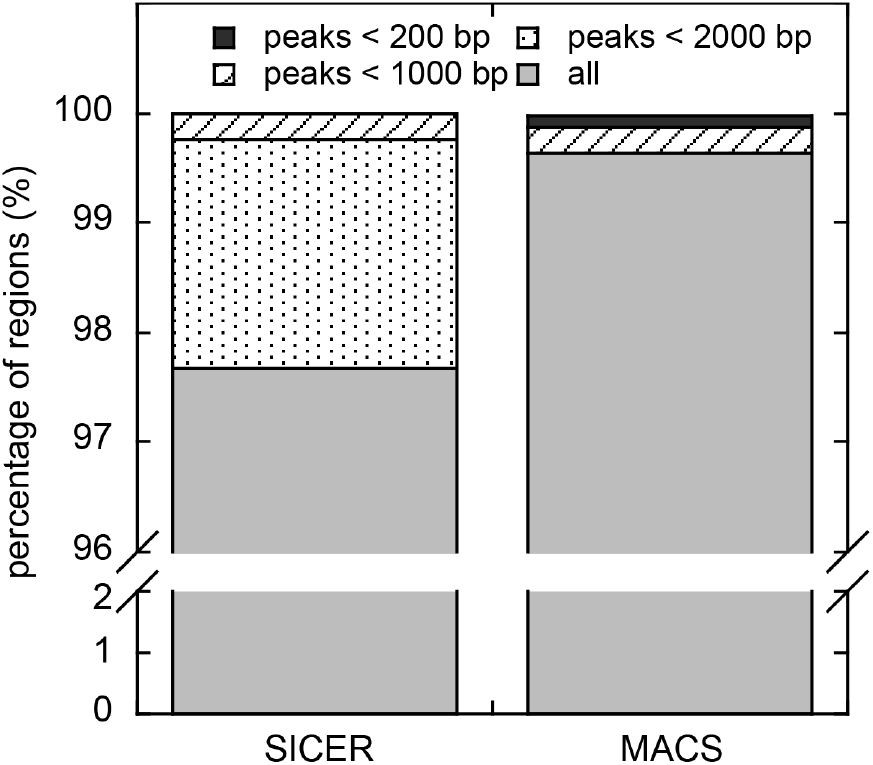
Peak calling summary for H3K9me3. MACS and SICER were used to identify peaks of H3K9me3 in NCs. Parameters were used as indicated in the Material and Methods section. Numbers of peaks with different sizes are given. 100% refers to all of the peaks identified by MACS (3630 peaks containing 0.4% of all mapped reads) or SICER (35780 peaks containing 9.45% of all mapped reads).

**Figure S13.**
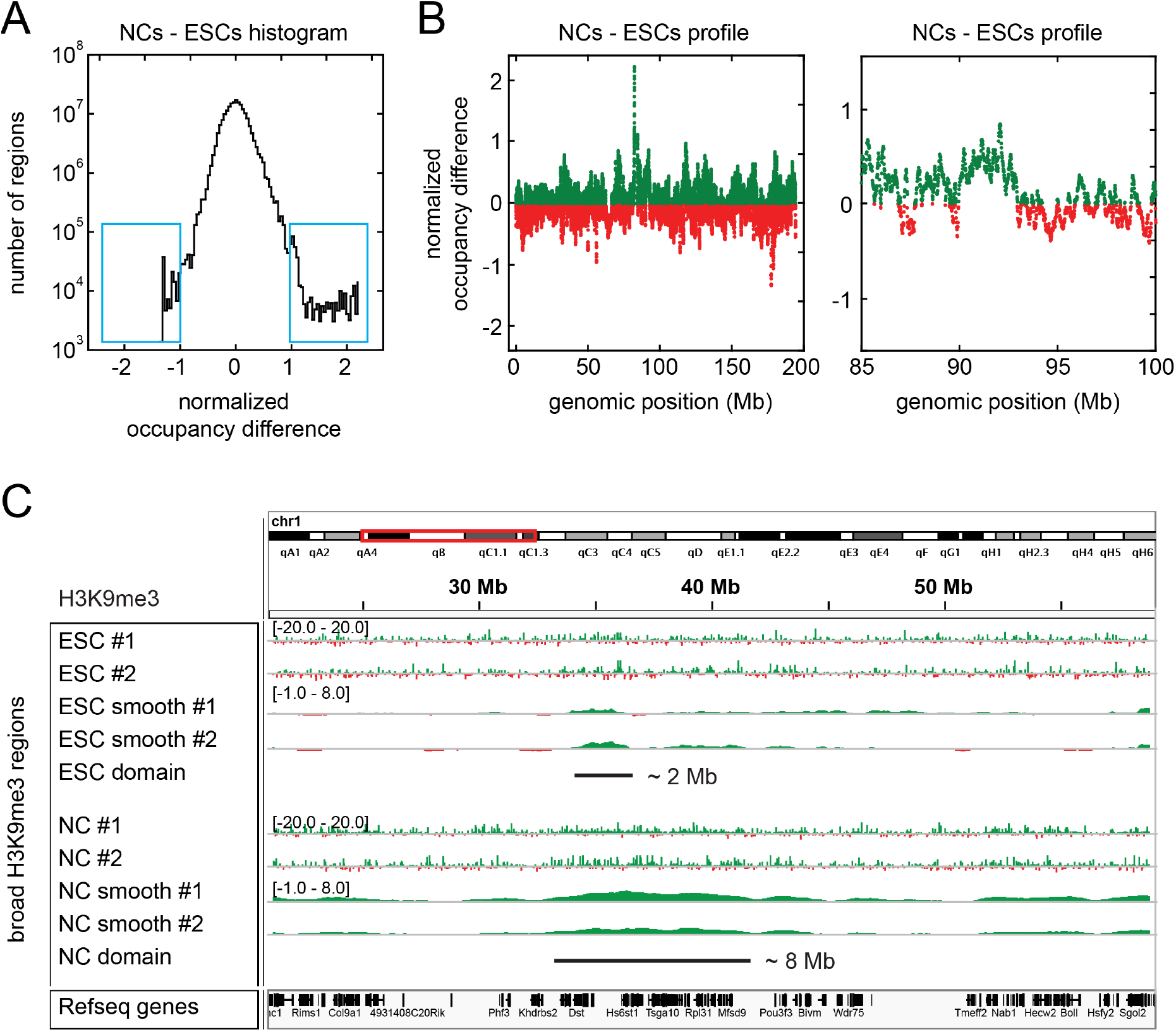
MCORE-directed annotation of chromatin features. MCORE identified broad H3K9me3 domains spanning on average 128 kb and 7.6 Mb in NCs. These domains were absent in ESCs, suggesting broadening of H3K9me3 domains during differentiation of ESCs into NCs (**Fig. 2**, *A* and *B* **Tables S3** and **S4**). **(A)** To identify broad regions enriched for H3K9me3 in NCs but to a lesser extent in ESCs, the coverage difference for normalized occupancy profiles in ESCs and NCs was calculated in a sliding window of 128 kb in size. A histogram for the obtained values is shown. The histogram is relatively symmetric and centered at zero, indicating that most genomic regions (that do not contain repetitive sequences) are not differentially modified with H3K9me3 in ESCs or NCs. The tails (blue rectangles) show that the largest coverage differences are found in regions that gain H3K9me3 in NCs. **(B)** The coverage difference along chromosome 1 (left, maximum and minimum values within 10 kb bins are plotted) and a zoom-in including the genomic region in **Fig. 2** *C* (88.7 - 89.3 Mb, right) are shown. **(C)**/b>To annotate the genomic positions of broad H3K9me3 domains, reads were counted and evaluated in a sliding window with the respective size. An example of a domain with ∼7.6 Mb that became broader in NCs is shown. For clarity the occupancy profiles were smoothed with 0.2-times the window size. An example for window size 128 kb is shown in **Fig. 2** *C*.

**Figure S14.**
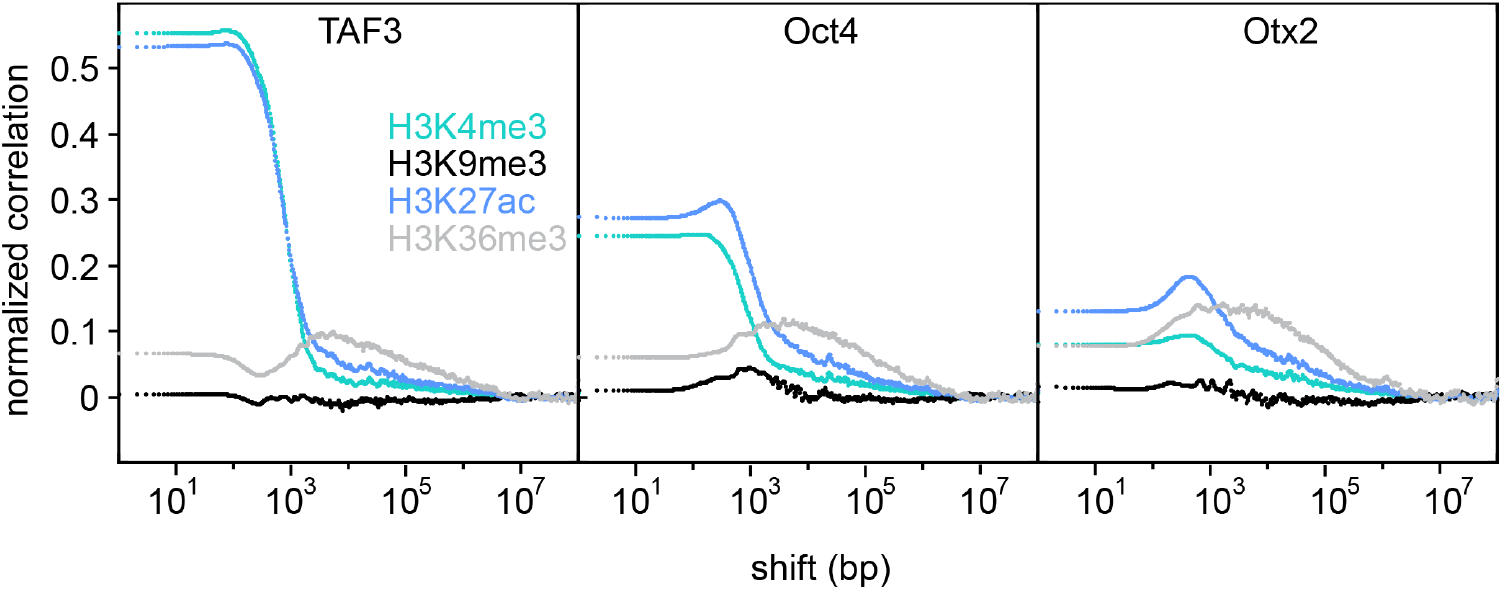
MCORE for transcription factor binding. Co-localization of transcription factors with different histone modifications was studied in ESCs. Cross-correlation functions of TAF3, Oct4 or Otx2 vs. H3K4me3, H3K9me3, H3K27ac and H3K36me3 are shown. Binding of TAF3 strongly correlates with H3K4me3 and H3K27ac, which mark active promoters and enhancers in mouse ESCs (1, 2). The binding of TAF3 to enhancers is in line with publications showing that active enhancers are transcribed by the RNA Polymerase II machinery (3) and that TAF3 mediates chromatin-looping events that regulate transcriptional activation (4). Oct4 and Otx2 are two transcription factors that regulate pluripotency and differentiation. Their binding correlates with H3K27ac in agreement with previous reports (5). The peaks in the correlation curves reflect the ∼300 bp distance between the binding site of the transcription factor and the modified nucleosome, which was also found recently (6). For each of the three transcription factors, maximum correlation with H3K36me3 was found at shift distances around 10 kb, which is similar to the average gene length and indicates that these factors globally bind adjacent to active genes. TAF3, Oct4 and Otx2 binding is uncorrelated with H3K9me3, which is consistent with their role in active transcription.

**Figure S15.**
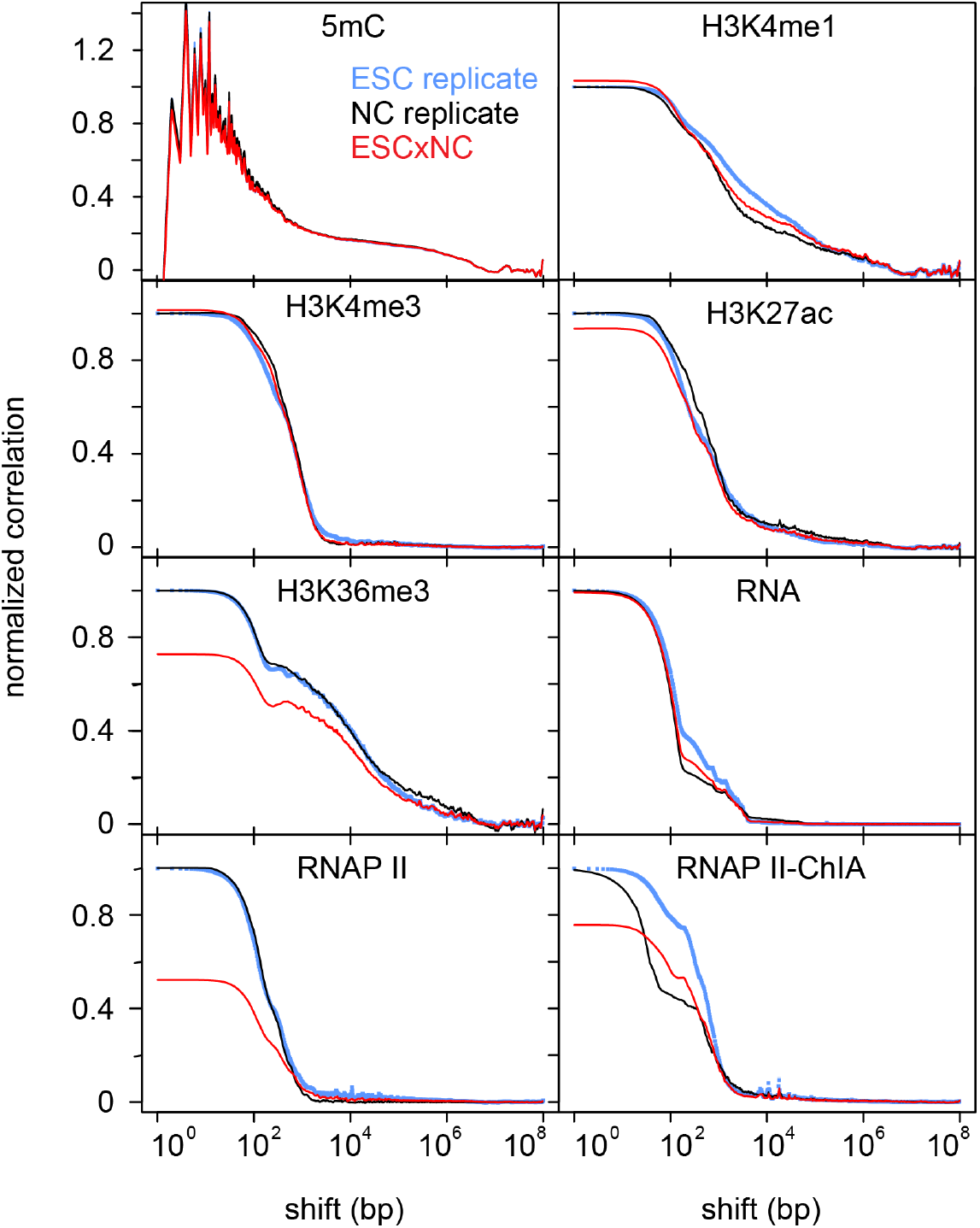
Spatial extension and co-localization of different features in ESCs versus NCs. Correlation functions for replicates of H3K4me1, H3K4me3, H3K27ac, H3K36me3 and RNA Polymerase II (RNAP II) ChIP-seq, RNA-seq (RNA) and RNAP II ChIA-PET data (RNAP II-ChIA) in ESCs (blue) and NCs (black) reflect the domain structures of the respective features. Cross-correlation functions (red) between the same feature in ESCs and NCs quantify the co-localization of this feature in both cell types. Most features depicted here did not drastically change their global distribution during differentiation because cross- and replicate correlation functions are similar to each other.

**Figure S16.**
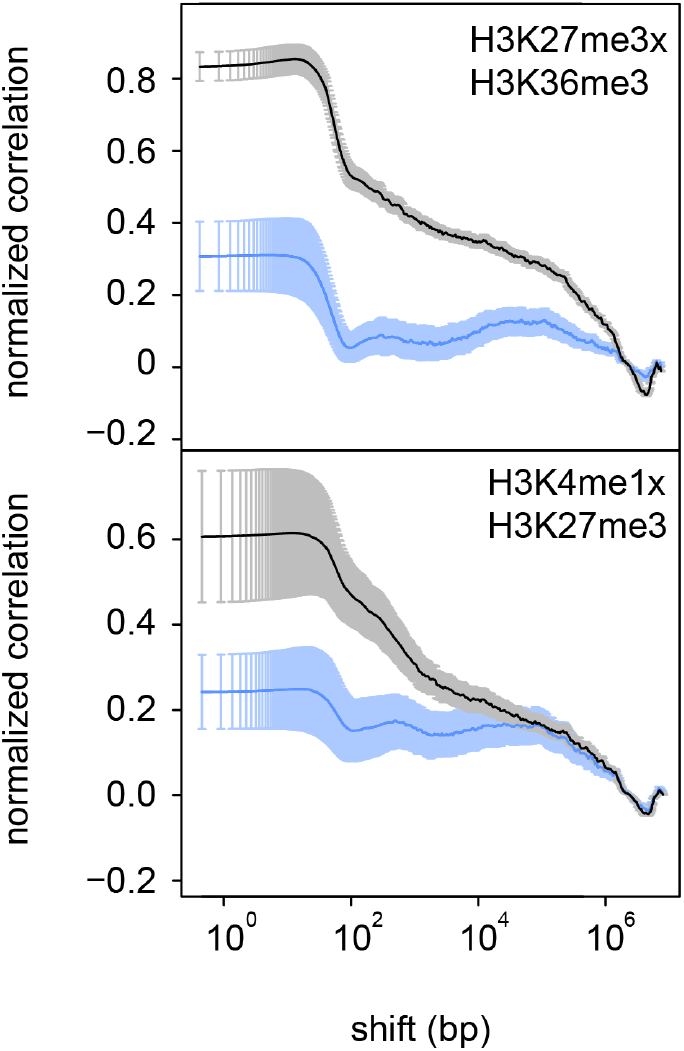
Heterochromatin reorganization during differentiation. Cross correlation functions between H3K27me3 and H3K4me1/H3K36me3 in ESCs (blue) or NCs (black) are shown. H3K27me3 exhibited increased co-localization with activating marks in NCs. Error bars indicate s.e.m. among replicates.

**Figure S17.**
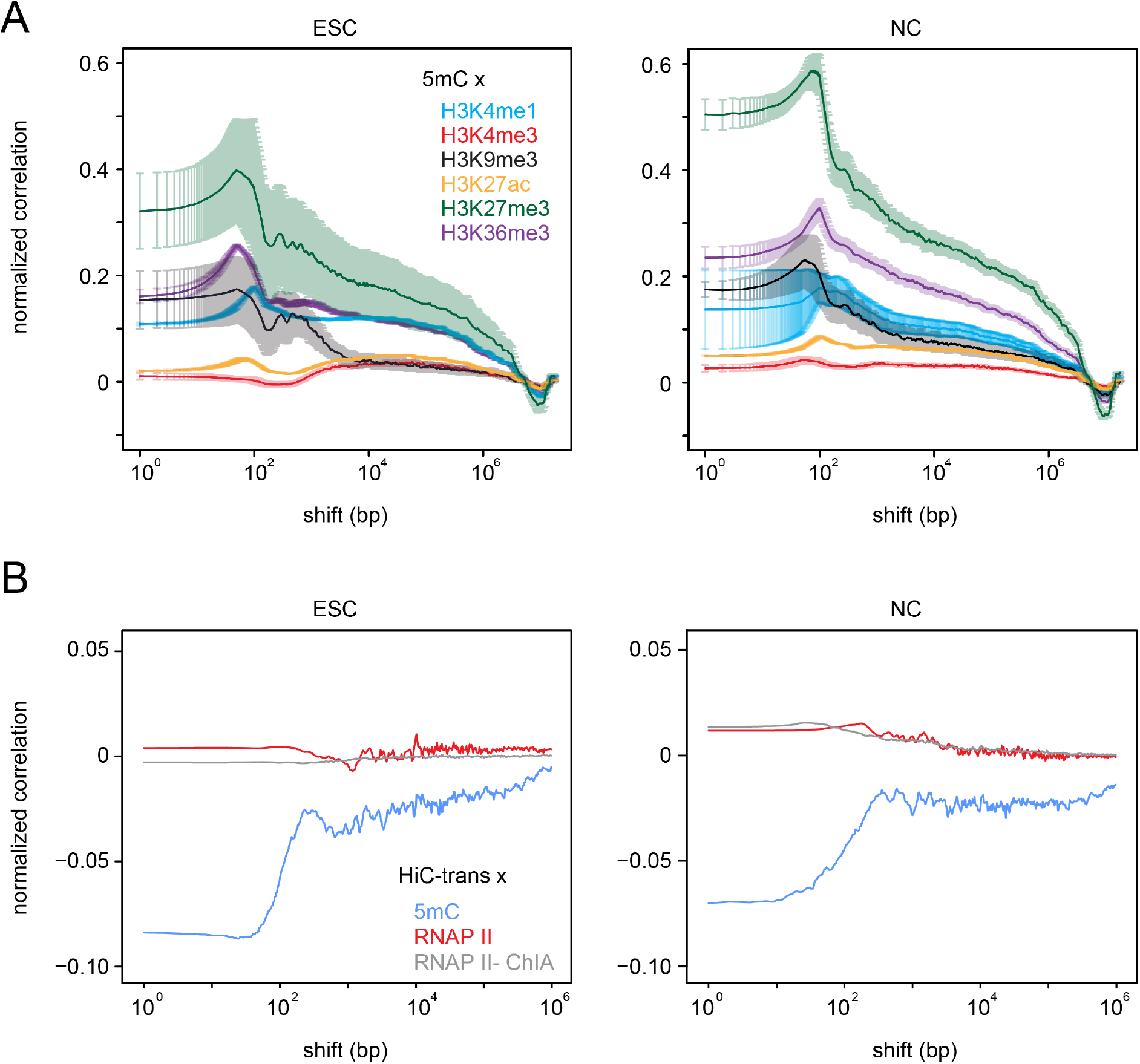
DNA methylation and inter-chromosomal contacts. **(A)** Cross correlation functions for DNA methylation and different histone modifications in ESCs (left) and NCs (right) are shown. Error bars indicate s.e.m. among replicates. **(B)** Cross-correlation functions for inter-chromosomal contact sites (Hi-C trans) and DNA methylation (5mC), RNA Polymerase II (RNAP II) and RNAP II ChIA-PET (RNAP II-ChIA) in ESCs (left) and NCs (right) are shown. RNAP II and RNAP II contact sites became moderately enriched at the surface of the chromosome territory in NCs, whereas 5mC tended to localize inside chromosome territories in both cell types. Small absolute values of correlation coefficients might be due to the relatively low number of inter-chromosomal contacts across the genome.

**Table S1.** Comparison of MCORE with other software tools.

|  | **Correlation function** | **Sliding window binning <sup>a</sup>** | **Peak <sup>a</sup> calling** | **Multi-scale representation** | **Probabilistic network models** | **Deconvolved correlation** | **Strand-specific correlation** |
| --- | --- | --- | --- | --- | --- | --- | --- |
| **Tool(s)** | MCORE | cisGenome, SiSSRs, SPP | MACS, SICER | MSR | ChromHMM, Segway | Arpeggio | SPP |
| **Platform** | Java | various | Python | Matlab script or compiler runtime | Java, Python | Java | R-script |
| **Sequencing data type** | Unrestricted | Unrestricted | ChIP-seq | Unrestricted | Unrestricted | ChIP-seq | ChIP-seq |
| **Mixed data type analysis implemented** | Yes | No | No | No <sup>b</sup> | Yes | No | No |
| **Applications** | Quality control, domain features, spatial relations | Local feature enrichment | Local feature enrichment | Multi-scale feature enrichment | Segmentation into feature states | Comparison of data sets, local structure | Quality control for sequencing data |
| **Correction <sup>c</sup>** | Input and/or control | Input or control | Input or control | Mappability, GC content, input or control | Input or control | Input or control | None |
| **Detected feature scale** | 1 bp – 1 chromosome | 1 bp – 1 chromosome | < 10 kb (MACS) variable (SICER) | 1 bp – 1 chromosome | 1 bp – 1 chromosome | 40 bp – 8 kb | Fixed window size |
| **Information on shifted relationships** | Yes | No | No | Limited <sup>b</sup> | No | No | No |
| **Required input parameters** | None | Window size | MACS: p-value threshold, tag length/shift<br>SICER: size of gap & window, FDR | Resolution, scale number, p-value threshold | State number, p-value threshold | None | None |
| **Number of data sets** | 2 | 1 | 1 | 1 <sup>b</sup> | >1 | 1 | 1 |
| **Noise sensitivity** | Low <sup>d</sup> | Low | High <sup>d</sup> | n.d. | n.d. | Low | Low |
| **Genome locus annotation** | No | Yes | Yes | Yes | Yes | No | Yes |
| **Output<sup>e</sup>** | Domain sizes, nucleosome spacing, spatial relationships, normalized occupancy | Enrichment over average | Local enrichment | Scale dependent enrichment | Length distribution, abundance of chromatin states | Feature profile, nucleosome spacing | Peak separation distance |
| **Operating system<sup>f</sup>** | All | All | All | Unix, Windows | All (ChromHMM)<br>Linux (Segway) | Unix, Mac OS X | All |
| **Comment** | Low sensitivity to noise, bias and undersampling | Read counting in a window of predefined size | Restricted scale-range | Can be applied as a peak caller with pruning. | Predefined number and type of states. | Removes large-scale structures by filtering | Recommended analysis prior to peak calling |
| **Reference** | This study | (7-9) | (10-12) | (13) | (14-17) | (18) | (7, 19) |

The table represents a non-comprehensive list of tools that are used to extract information about chromatin features from deep sequencing data sets.
<sup>a</sup> Exemplary tools are mentioned. For other programs see compilations in ref. (20, 21).
<sup>b</sup> MSR can be applied to identify region of simultaneous enrichments for two different ChIP-seq data sets by computing a matrix of segments, but this analysis is not part of the current implementation. In some cases differential correlation of the matrix indicates the presence of shifted correlations.
<sup>c</sup> Control reactions depend on the type of sequencing data and could involve for example a ChIP-seq reaction without the specific antibody.
<sup>d</sup> See ref. (22) for peak calling and **Figs. S5** and **S6** for MCORE
<sup>e</sup> The “enrichment” analysis of a given feature would also provide the information about its depletion with respect to a given average signal.
<sup>f</sup> All operating systems refers to Unix, Windows and Mac OS X.

**Table S2.** Overview of chromatin features assessed in this study. Due to the plethora of functions associated with each feature only a coarse-grained assignment of the most important function is provided.

| Feature | Function | Location | Reference |
| --- | --- | --- | --- |
| 5mC | repression, splicing, TF binding | CpG dinucleotides | (23) |
| H3K4me1 | poised | enhancer | (24) |
| H3K4me3 | activation | promoter | (25) |
| H3K9ac | activation | promoter | (26) |
| H3K9me3 | repression | promoter, enhancer, repeats | (27) |
| H3K27ac | activation | promoter, enhancer | (24) |
| H3K27me3 | repression | promoter, enhancer, CpG islands | (27) |
| H3K36me3 | activation, splicing | active gene bodies | (28, 29) |
| H3K4me1,<br>H3K27ac | activation | enhancer | (24) |
| H3K4me3,<br>H3K27me3 | bivalent | promoter | (30) |
| RNAP II | transcription | promoter, active gene bodies,<br>active nuclear compartments | (31, 32) |
| RNAP II<br>ChIA-PET | promoter-promoter/<br>enhancer interactions | promoter, enhancer | (33, 34) |
| Hi-C trans | interactions between two<br>chromosomes | surface of<br>chromosome territory | (35, 36) |
| RNA | transcript | transcribed chromatin | (37) |

**Table S3.** Fit parameters for selected correlation functions in ESCs. Correlation functions calculated for replicates of H3K4me3, H3K9me3, H3K27me3 and H3K36me3 (**Figs. 2** *A* and **S11**) were fitted with Eq. 7 (Materials and Methods), yielding the indicated fit parameters and corresponding standard errors (SE). The minimum number of domains required to yield uncorrelated fit residuals was chosen. The amplitudes a1-a5 represent the relative domain abundance, the decay length parameters b1-b5 represent half of the respective domain sizes, and the value of c2 reflects nucleosome spacing. See text and Materials and Methods for further details.

| ESC | H3K4me3 |  | H3K9me3 |  | H3K27me3 |  | H3K36me3 |  |
| --- | --- | --- | --- | --- | --- | --- | --- | --- |
| number of domains | 3 |  | 4 |  | 5 |  | 3 |  |
|  | value | SE | value | SE | value | SE | value | SE |
| a1 (%) | 18.0 | 0.5 | 27.6 | 0.6 | 25.3 | 1.2 | 26.9 | <0.5 |
| a2 (%) | 75.7 | 0.6 | 46.4 | 2.4 | 20.0 | 4.3 | 51.4 | 3.0 |
| a3 (%) | 6.3 | 0.3 | 21.0 | 3.0 | 23.3 | 6.0 | 21.7 | 1.8 |
| a4 (%) | - | - | 5.0 | 3.9 | 22.1 | 3.5 | - | - |
| a5 (%) | - | - | - | - | 9.3 | 8.3 | - | - |
| b1 (bp) | 132 | 2 | 107 | 2 | 106 | 3 | 119 | 2 |
| b2 (bp) | 926 | 6 | 1586 | 18 | 3198 | 173 | 14803 | 296 |
| b3 (kb) | 33 | 6 | 11 | 2 | 16 | 2 | 356 | 105 |
| b4 (kb) | - | - | 1121 | 704 | 322 | 46 | - | - |
| b5 (kb) | - | - | - | - | 4481 | 195 | - | - |
| c1 (%) | 99 | fixed | 98 | <0.05 | 69 | 1 | 97 | 1 |
| c2 (bp) | 173 | fixed | 182 | 9 | 207 | 5 | 182 | 5 |
| c3 (bp) | 1000 | fixed | 654 | 340 | 219 | 9 | 802 | 303 |
| n1 | 1.97 | 0.05 | 2.20 | 0.10 | 3.31 | 0.27 | 2.30 | 0.50 |
| n2 | 1.25 | 0.01 | 1.11 | 0.00 | 1.96 | 0.25 | 0.62 | 0.01 |
| n3 | 0.38 | 0.02 | 0.64 | 0.10 | 1.28 | 0.33 | 0.45 | 0.04 |
| n4 | - | - | 0.39 | 0.10 | 0.79 | 0.17 | - | - |
| n5 | - | - | - | - | 3.96 | 0.97 | - | - |

**Table S4.** Fit parameters for selected correlation functions in NCs. Correlation functions calculated for replicates of H3K4me3, H3K9me3, H3K27me3 and H3K36me3
(**Figs. 2 B** and *S11*) were fitted with Eq. 7, yielding the indicated fit parameters and corresponding standard errors (SE) as described in the Materials and Methods section and the legend to Table S3.

| NC | H3K4me3 |  | H3K9me3 |  | H3K27me3 |  | H3K36me3 |  |
| --- | --- | --- | --- | --- | --- | --- | --- | --- |
| number of domains | 3 |  | 4 |  | 4 |  | 3 |  |
|  | value | SE | value | SE | value | SE | value | SE |
| a1 (%) | 18.2 | 1.5 | 47.5 | 3.1 | 54.7 | 1.2 | 25.7 | 0.4 |
| a2 (%) | 79.9 | 1.5 | 23.2 | 3.6 | 11.5 | 1.9 | 57.3 | 0.9 |
| a3 (%) | 1.9 | 1.9 | 17.7 | 2.9 | 17.5 | 2.4 | 17.0 | 0.9 |
| a4 (%) | - | - | 11.6 | 5.5 | 16.3 | 3.2 | - | - |
| b1 (bp) | 243 | 4 | 202 | 19 | 111 | 2 | 111 | 1 |
| b2 (bp) | 985 | 14 | 2036 | 142 | 1791 | 91 | 11848 | 287 |
| b3 (kb) | 617 | 106 | 64 | 13 | 48 | 9 | 1412 | 85 |
| b4 (kb) | - | - | 3771 | 256 | 3132 | 131 | - | - |
| c1 (%) | 98 | <0.5 | 74 | 4 | 82 | 2 | 99 | <0.5 |
| c2 (bp) | 134 | 3 | 175 | 4 | 218 | 15 | 182 | 6 |
| c3 (bp) | 3017 | 3017<sup>a</sup> | 367 | 41 | 224 | 27 | 11505 | 11505<sup>a</sup> |
| n1 | 2.01 | 0.12 | 2.31 | 0.60 | 2.67 | 0.09 | 2.47 | 0.07 |
| n2 | 1.43 | 0.03 | 1.11 | 0.15 | 1.54 | 0.24 | 0.59 | 0.01 |
| n3 | 0.53 | 0.07 | 0.62 | 0.13 | 0.52 | 0.09 | 0.79 | 0.05 |
| n4 | - | - | 1.56 | 0.24 | 1.64 | 0.15 | - | - |

**Table S5.** Summary of data sets used in this study.

| target | cell type | accession<br>replicate1 | accession<br>replicate2 | reference |
| --- | --- | --- | --- | --- |
| Input | ESC | GSM1516068 | GSM1516069 | This study |
| Input | ESC | SRX499123 | SRX499124 | (5) |
| IgG | ESC | GSM1516070 | GSM1516071 | This study (RA073) |
| IgG | ESC | GSM1516072 | GSM1516073 | This study (PP500P) |
| IgG | ESC | SRR331056 | SRR331057 | (4) |
| 5mC | ESC | SRX080191 |  | (38) |
| H3K27ac | ESC | GSM1516076 | GSM1516077 | This study (ab4729) |
| H3K27me3 | ESC | GSM1516074 | GSM1516075 | This study (ab6002)) |
| H3K36me3 | ESC | GSM1516082 | GSM1516083 | This study (ab9050) |
| H3K4me1 | ESC | GSM1516080 | GSM1516081 | This study (ab8895) |
| H3K4me3 | ESC | GSM1516086 | GSM1516087 | This study (ab8580) |
| H3K9me3 | ESC | GSM1516084 | GSM1516085 | This study (ab8898) |
| Hi-C | ESC | SRX116341 | SRX116342 | (35) |
| Input | ESC | SRR317225 | SRR317226 | ENCODE |
| Oct4 | ESC | SRX499114 | SRX499115 | (5) |
| Otx2 | ESC | SRX499116 | SRX499117 | (5) |
| RNA | ESC | GSM1516088<br>GSM1516089 | GSM1516090<br>GSM1516091 | This study |
| RNAP II | ESC | SRR489721 | SRR489722 | ENCODE |
| RNAP II-ChIA | ESC | SRX243706 | SRX243707 | (34) |
| TAF3 | ESC | SRR331054 | SRR331055 | (4) |
| Input | NPC | SRX604258 | SRX604259 | (39) |
| IgG | NPC | GSM1516092 | GSM1516093 | This study (RA073) |
| 5mC | NPC | SRX080193-5 |  | (38) |
| H3K27ac | NPC | GSM1516096 | GSM1516097 | This study (ab4729) |
| H3K27me3 | NPC | GSM1516094 | GSM1516095 | This study (ab6002)) |
| H3K36me3 | NPC | SRX604262 | SRX604263 | (39) |
| H3K4me1 | NPC | GSM1516100 | GSM1516101 | This study (ab8895) |
| H3K4me3 | NPC | GSM1516102 | GSM1516103 | This study (ab8580) |
| H3K9me3 | NPC | SRX604260 | SRX604261 | (39) |
| Hi-C | Cortex | SRX128219 | SRX128220 | (35) |
| Input | Brain E14.5 | SRR489727 | SRR578284 | ENCODE |
| RNA | NPC | GSM1516104<br>GSM1516105 | GSM1516106<br>GSM1516107 | This study |
| RNAP II | Brain E14.5 | SRR578272 | SRR578273 | ENCODE |
| RNAP II-ChIA | NPC | SRX243710 |  | (34) |

